# Interpretable modeling of genotype-phenotype landscapes with state-of-the-art predictive power

**DOI:** 10.1101/2021.06.11.448129

**Authors:** Peter D. Tonner, Abe Pressman, David Ross

## Abstract

Large-scale measurements linking genetic background to biological function have driven a need for models that can incorporate these data for reliable predictions and insight into the underlying biochemical system. Recent modeling efforts, however, prioritize predictive accuracy at the expense of model interpretability. Here, we present LANTERN (https://github.com/usnistgov/lantern), a hierarchical Bayesian model that distills genotype-phenotype landscape (GPL) measurements into a low-dimensional feature-space that represents the fundamental biological mechanisms of the system while also enabling straightforward, explainable predictions. Across a benchmark of large-scale datasets, LANTERN equals or outperforms all alternative approaches, including deep neural networks. LANTERN furthermore extracts useful insights into the landscape including its inherent dimensionality, a latent space of additive mutational effects, and novel metrics of landscape structure. LANTERN facilitates straightforward discovery of fundamental mechanisms in GPLs, while also reliably extrapolating to unexplored regions of genotypic-space.

## 1 Introduction

Genotype-phenotype landscapes (GPLs) characterize the relationship between a gene’s mutations and its function. Driven by reduced sequencing costs and growing experimental throughput, the scale of available GPL measurements has increased dramatically, with recent measurements sampling as many as 10^4^–10^7^ distinct genotypes (Kinney et al. 2019). These measurements play an expanding role in understanding biological variation, with applications from engineering to epidemiology (Tack et al. 2021; Starr, Greaney, et al. 2020). Despite their importance, though, even the highest-throughput experimental measurements cannot overcome the massive combinatorial size of GPLs. For example, meaningful changes in engineered function or virulence are often the result of three or more mutations. For a typical protein, there are approximately 10^11^ potential triple mutants, so a large-scale GPL measurement on the order of 10^4^–10^7^ observations can only sample a tiny fraction of the relevant mutational space around the native sequence. Since this makes complete experimental coverage of GPLs unrealistic, a full understanding of landscape spaces can only come from estimates of un-measured genotype-phenotype combinations, using models and their predictions.

Predicting phenotypes for genotypes with multiple mutations is challenging because the effect of each mutation depends on which other mutations are present. So, the phenotypic change due to multiple mutations is not simply the sum of the changes from each single mutation (Domingo et al. 2019). This context dependence, referred to as *epistasis*, arises from many biological mechanisms and has motivated diverse modeling approaches. For example, specific pairs of protein residues can have strong non-additive interactions due to their close physical proximity in the fold-protein, and GPL models can represent this effect through pair-wise interaction coefficients (Poelwijk et al. 2016). Alternatively, even when individual mutations cause additive changes to an underlying biophysical parameter (e.g. folding free energy), the measured phenotype can be a non-linear function of that parameter (Jakub Otwinowski et al. 2018; Jakub Otwinowski 2018; Starr and Thornton 2016). GPL models that directly represent these epistatic mechanisms have the advantage of being *interpretable* (Lipton 2018). Interpretability comes from the correspondence between model components and biological mechanisms and can provide practitioners with an intuitive understanding of each component of the larger model. Additionally, clear explanations of how the model generates each prediction increases interpretability by aiding the diagnosis of innacurate predictions, increasing the reliability of decisions made from predictions, and increasing trust in the model.

Existing interpretable GPL models often have limitations on their predictive accuracy (J. Otwinowski et al. 2014). So, recent efforts to model large-scale GPL measurements have instead relied on deep neural network (DNN) architectures due to their superior predictive performance (Pokusaeva et al. 2019; Sarkisyan et al. 2016; Valeri et al. 2020; Angenent-Mari et al. 2020; Ching et al. 2018). DNNs make predictions through complicated cascades of non-linear computation with large numbers of parameters that lack direct biological motivation, operating essentially as a black-box. To make DNNs more interpretable, post hoc *explainability* techniques estimate the relevant factors of DNN predictions (Jiménez-Luna et al. 2020). By necessity, however, these methods only approximate the actual important features and incorrect conclusions can be drawn by mischaracterization of the details in the explanation (Rudin 2019). Additionally, these techniques only apply to individual predictions rather than generalizing the whole model (Guidotti et al. 2019). So, explaining these black-box models does not straightforwardly scale to the billions of predictions that may be of interest for GPL-dependent research. Overall, DNNs provide a useful approach for maximizing predictive accuracy but force users to make compromises on model interpretability.

Here, we address the conflict between predictive accuracy and interpretability by developing a novel GPL modeling approach called LANTERN (a genotype-phenotype <u>lan</u>dscape in<u>ter</u>pretable <u>n</u>onparametric model). LANTERN learns interpretable models of GPLs by finding a latent, low-dimensional space where mutational effects combine *additively*. LANTERN then captures the non-linear effects of epistasis through a multi-dimensional, non-parametric Gaussian-process model. In a benchmark across multiple protein GPLs, LANTERN achieves predictive accuracy as good as or better than existing models, including DNNs. Importantly, LANTERN automatically provides interpretable explanations of these predictions. LANTERN therefore remains highly interpretable while maximizing predictive power, and can thus increase the coverage of GPLs by orders of magnitude while simultaneously distilling complex landscapes into their fundamental structure.

## 2 Results

### 2.1 Constructing interpretable models of genotype-phenotype landscapes

LANTERN takes as input data a combination of genotypes and their measured phenotypes (Fig 1a). A LANTERN model of this data has two key components. First, LANTERN decomposes genetic mutations onto a latent mutational effect space, where individual mutations are each represented by a vector (e.g. **z**^(*i*)^ for mutation *i*, Fig 1c). Vectors provide interpretable comparisons between mutations, with two vectors in the same direction implying similar function and vector magnitude representing strength of effect. Importantly, we assume that individual mutations combine additively, represented as vector addition in this latent mutational effect space. This underlying additive structure can accurately represent many biophysical phenomena. For example, individual mutations often additively impact the thermodynamic stability of protein folding (Tokuriki et al. 2009). GPLs measurements, however, frequently target a phenotype that is a non-linear function of these additive effects. In the example of thermodynamic folding stability, many protein phenotypes remain robust to small decreases in this stability but rapidly diminish beyond a certain threshold (Starr and Thornton 2016). So, the second component of LANTERN relates measured phenotypes to the combination of latent mutational effects through a smooth, non-linear surface: *f* (**z**)(Fig 1d).

**Figure 1:**
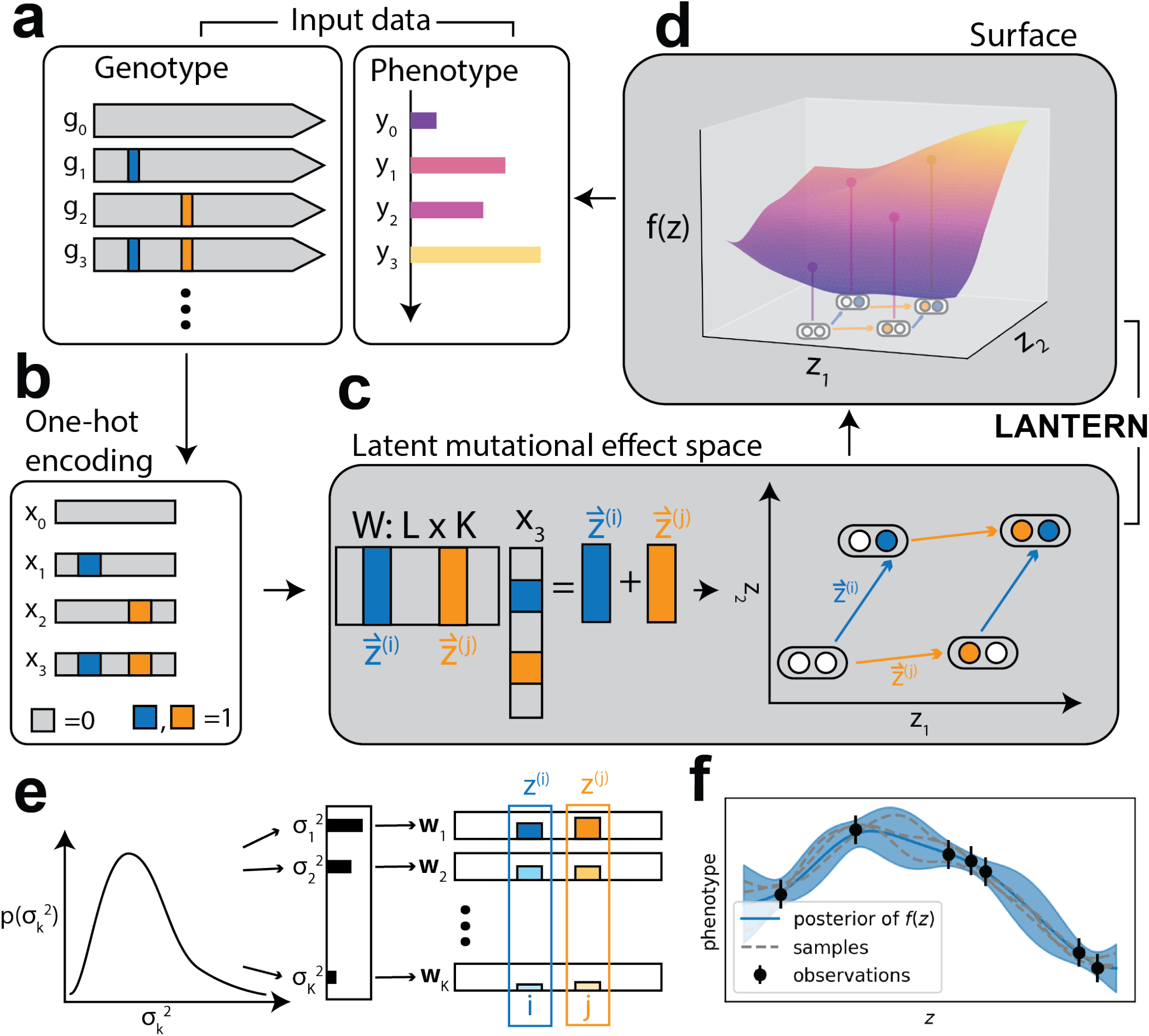
Interpretable modeling of genotype-phenotype landscapes. (a) LANTERN takes as input genotype-phenotype measurements, where each genetic background (genotype) has a corresponding measured phenotype. (b) LANTERN converts the genotype of each variant ***k*** into a one-hot encoded vector ***x***_***k***_ ϵ {0, 1} ^***p***^, where ***p*** is the total number of mutations observed across all variants and ***x***_***kl***_ = 1 implies the presence of mutation ***l*** in variant ***k*** (*x*_***kl***_ = 0 otherwise) (c) LANTERN predicts the position of variant ***k*** in the latent mutational effect space as a linear combination of mutation effect vectors with an unknown matrix *W* to be learned from the data. This formulation represents the assumption that mutations combine *additively* in the latent space. Additionally, individual mutations have an interpretable representation in the model in the form of their mutational effect vector 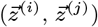. Computing the location of each variant *k* in the latent mutational effect space then requires simply adding the mutational effect vectors for mutations present in the variant. (d) Observed phenotypes are a non-linear function *f* (***z***) of the latent mutational effect space, which ultimately results in a non-linear relationship between genotype and phenotype. (e) Dimensionality of the model is controlled through a hierarchical prior on latent mutational effect dimensions. The variance of individual dimensions in the latent space (e.g. the rows of ***W***) have a prior skewed to small values. This results in the majority of dimensions effectively shrinking towards zero, and the smallest dimensionality necessary to explain the data is recovered. (f) Non-linear functions *f* (***z***) are modeled with Gaussian process priors. An example GP posterior is shown, fit to observations (black dots, bars show observation uncertainty), where the solid blue line is the posterior predictive mean and the shaded region is the 95% credible region. Dotted lines show example function draws from this posterior.

These two components, the latent mutational effect space **z** and non-linear surface *f* (**z**), make LANTERN interpretable. Both components clearly correspond to general biophysical mechanisms often seen in GPL measurements, and therefore have intuitive explanations for their role in the model. Additionally, they explain the predictions made by LANTERN straightforwardly: each prediction results by first combining the latent mutational effects and then transforming through the surface *f* (**z**). Notably, LANTERN makes no assumptions about the specific biochemical mechanisms driving the complexity of GPLs in either the latent mutational space or the non-linear response surface. Instead, LANTERN learns these relationships directly from the GPL data after assuming the general structure outlined above. Therefore, while each component learned by LANTERN may have a direct correspondence to the biophysical mechanisms of the systems, these connections must be determined through additional analysis.

For application to GPL data, we implemented LANTERN as a hierarchical Bayesian model. As part of this hierarchy, we employed an approach for determining the dimensionality of any GPL directly from the data. Specifically, we learn a relative scale between each dimension of the latent mutational effect space (**z**) in the form of each dimension’s variance. The variance of each dimension determines its relative impact on describing the effect of mutations: dimensions with higher variance result in larger mutational effects while dimensions with variance close to zero are effectively removed from the model (Bishop 1999). The variance of each dimension therefore reflects its relative impact on the model, which we refer to as the *relevance* of the dimension. To ensure that LANTERN learns the minimal number of dimensions necessary to explain the data, we placed a hierarchical prior on the variance of each dimension of the latent mutational effect space (Fig 1d). This prior represents the expectation that only a relatively small number of latent mutational effect dimensions will be necessary to explain the data and ensures that dimensions that are not supported by the data are removed. LANTERN therefore learns the minimal number of dimensions necessary to explain the data, which we refer to as the *dimensionality* of the GPL.

To learn the non-linear relationship between latent mutational effects and measured phenotypes, we placed a Gaussian process (GP) prior on the surface *f* (**z**) (Fig 1f) (Rasmussen et al. 2006). GPs recover the distribution of possible functions that best explain the data, rather than specifying a parametric form to the underlying relationship between *z* and the observed phenotypes governed by *f* (Supplemental Fig S1). This ensures that LANTERN can learn the surface *f* (**z**) of *any* GPL automatically from the data rather than relying on expert knowledge to chose an appropriate parametric form (Jakub Otwinowski et al. 2018; Sailer, Shafik, et al. 2020). To learn both components of LANTERN for different GPLs, we apply stochastic variational methods that make inference tractable and scalable to millions of observations (Hoffman et al. 2013).

### 2.2 LANTERN learns biophysical mechanisms

To determine how well LANTERN can discover true biophysical mechanisms from GPL data, we evaluated its performance on simulated data from an analytic model of protein allostery (Razo-Mejia et al. 2018). Allosteric proteins regulate cellular processes in response to changes in ligand concentration, and the analytic model describes this response as a function of the underlying biophysical constants such as ligand-protein binding constants and free-energy differences between protein states. In the analytic model, the biophysical constants determine three phenotypic parameters that characterize the allosteric dose-response curve: the basal and saturating transcription levels (G_0_ and G_∞_, respectively), and the concentration of ligand where transcription is halfway between minimum and maximum (EC_50_).

Briefly, we constructed synthetic GPL measurements by simulating a set of mutations that additively shift the wild-type biophysical constants (Fig 2a, Section 4.6). We then randomly combined these mutations to simulate individual protein variants with a mean of 4.38 mutations per variant. For each variant, the resulting perturbed biophysical constants determined the allosteric dose-response parameters via the analytic model. We trained LANTERN only using the variant genotypes and resulting dose-response parameters, *without* access to the underlying biophysical parameters or the analytical model.

**Figure 2:**
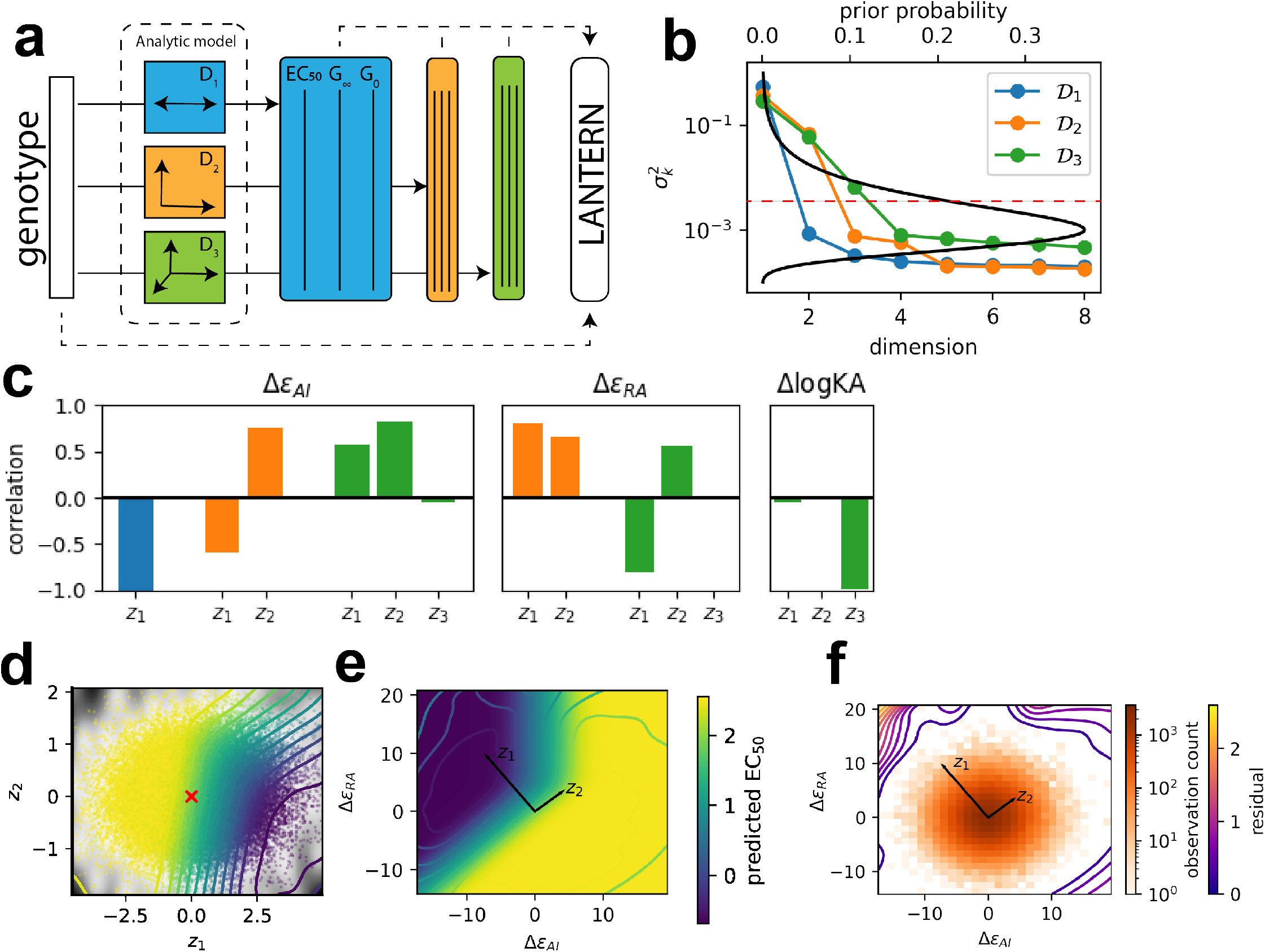
Recovering biophysical landscapes and dimensions. (a) Simulating allosteric GPL landscapes of varying dimensionality. Randomized genotypes have latent mutational effects on one to three underlying biophysical constants (𝒟 _1_ to 𝒟_3_). The analytical model determines the allosteric parameters EC_50_, G∞, and 𝒢_0_ for each resulting combinations of biophysical constants. LANTERN learns a model for each dataset, without knowledge of the underlying biophysics. (b) Approximate posterior variance of independent dimensions for each allosteric simulations of varying latent dimensionality. Black line shows the prior distribution. To determine the dimensionality learned by LANTERN on each simulated dataset, we counted the number of dimensions with posterior mean variance with less than ∼0.2% prior probability (red line). (c) Correlation between learned latent dimensions and simulated biophysical parameters. (d) Learned two-dimensional surface *f* (**z**) for EC_50_ in 𝒟_2_. Contours show the posterior mean of *f* (**z**), shading shows the relative variance of *f* (**z**) and scatter points are observations positioned by their latent **z** value and colored by their observed parameter value. (e) Rotation of the learned surface *f* (**z**) in (d) to the true biophysical surface. As a function of biophysical parameters Δ*∈*_***AI***_ and Δ*∈*_***RA***_, the image shows the true biophysical landscape and the contours show the posterior mean of *f* (**z**) from LANTERN. The black vectors mark the rotation of the **z**_1_ and **z**_2_ dimensions to the native biophysical space. (f) Squared residual between posterior mean of *f* (**z**) and true EC_50_ surface shown as contours rotated as in (e) overlayed on the sampling density of variants in the latent space. See section 4.6 for method details.

To test LANTERN’s ability to correctly identify GPL dimensionality, we simulated three different GPL datasets. In these datasets, we varied the number of perturbed biophysical constants from one to three (datasets 𝒟_1_ to 𝒟_3_, respectively), creating datasets with true dimensionality equal to the number of varying biophysical constants. For each dataset, we considered the learned variance of the latent-space dimensions (Fig 2b) and determined the dimensionality learned by LANTERN by counting the number of dimensions with learned variance above a fixed threshold. Below this threshold, no latent mutational effects remain significantly non-zero (Supplemental Fig S2a). Additionally, when we removed dimensions with variance below this threshold from the model, predictive accuracy remained constant (Supplemental Fig S2b–f). Across all simulated datasets, the learned dimensionality exactly matched the number of biophysical constants perturbed when generating each landscape, demonstrating that LANTERN can recover the true dimensionality of GPL measurements automatically from data.

To determine if LANTERN also learns representations of GPLs that agree with the underlying biophysical mechanisms, we compared the model learned by LANTERN to the true biophysical system. LANTERN learned latent mutational effects that strongly correlated with true biophysical parameters, despite the fact that LANTERN had no access to this information (Fig 2c, Supplemental Fig S3). Furthermore, LANTERN learned smooth, interpretable surfaces that accurately predict the allosteric phenotype of each simulated landscape (Fig 2d, Supplemental Figs S4ab,S5). Using the correlation between latent mutational effects learned by LANTERN and the true biophysical constants in the simulation (Fig 2c), we rotated and re-scaled the space of latent mutational effects to directly compare *f* (**z**) to the true biophysical surface (Fig 2e, Supplemental Fig S4c). In regions of latent mutational effect space with large sampling coverage, the rotated and re-scaled surface, *f* (**z**), matches the surface from the analytical model exactly (Fig 2e). As sampling density decreases, the deviation between predicted and true surfaces increases (Fig 2f). LANTERN therefore approximates a rotated and scaled version of the true biophysical model in regions with high experimental coverage, but balances this against uncertainty in more sparsely sampled regions of parameter space. Overall, this demonstrates that LANTERN can recover the underlying dimensionality and biophysical mechanisms of our simulations *de-novo* from data, without additional domain specific knowledge.

### 2.3 LANTERN outperforms alternative predictive methods

To compare LANTERN’s predictive accuracy to alternative models with real GPL data, we analyzed three published large-scale GPL datasets (Table 1): the fluorescence brightness of green fluorescent protein (**avGFP**; Sarkisyan et al. 2016), the dose-response curves of an allosteric transcription factor (**LacI**; Tack et al. 2021), and the joint phenotypes of ACE2 binding affinity and structural stability for the receptor binding domain of the SARS-CoV-2 spike protein (**SARS-CoV-2**; Starr, Greaney, et al. 2020). Each dataset samples a large number of genotypes, between ∼50,000 and ∼170,000 distinct genotypes and ∼1,800 to ∼4,000 unique mutations, making them ideal candidates for evaluating predictive performance of different GPL models. We evaluated GPL model performance on each dataset through predictive accuracy (*R*^2^) cross-validated over ten random splits of the data into test and training sets.

**Table 1:** Large-scale genotype phenotype landscape datasets.

| Dataset | observations | mutations | phenotype | reference |
| --- | --- | --- | --- | --- |
| avGFP | 54,025 | 1,879 | fluorescence | (Sarkisyan et al. 2016) |
| LacI | 47,462 | 2,510 | hill coefficient $EC_{50}$ | (Tack et al. 2021) |
| LacI | 47,462 | 2,510 | hill coefficient $G_{\infty}$ | (Tack et al. 2021) |
| SARS-CoV-2 | 146,437 | 3,802 | ACE2-RBD binding ($\log K_D$) | (Starr, Greaney, et al. 2020) |
| SARS-CoV-2 | 177,759 | 4,002 | RBD expression ($\Delta \log \text{MFI}$) | (Starr, Greaney, et al. 2020) |

To provide the broadest comparison of LANTERN to alternatives, we tested multiple modeling approaches with each dataset, including interpretable linear and non-linear models as well as black-box DNNs. Overall, LANTERN achieves a new state-of-the-art in predictive accuracy. In all but one instance, LANTERN equalled or outperformed all alternative approaches with respect to ten-fold cross-validated prediction accuracy (see Methods, Fig 3a). For avGFP, all of the tested models except for a simple linear approach have comparable predictive accuracy. In this case, simpler models than LANTERN are sufficient to make accurate predictions. Importantly, LANTERN achieves the same predictive accuracy as these approaches, meaning that LANTERN does not overfit the data despite its capacity to learn more complex models. For SARS-CoV-2, LANTERN has higher predictive accuracy than all alternatives other than a dense feedforward neural network, which performed equally well. Additionally, LANTERN outperforms all other approaches in predicting LacI EC_50_. In one case, the G_∞_ of LacI, a one-dimensional DNN predicts out-of-sample phenotypes more accurately than LANTERN. The G_∞_ phenotype poses unique challenges due to the complex relationship between observed G_∞_ and phenotype uncertainty: values of G_∞_ most distinct from the wild-type are also the most uncertain (Tack et al. 2021). This may explain the slight decrease in predictive accuracy for LANTERN in this case. Notably, when adapting the variational loss to more robustly handle highly uncertain measurements (Jankowiak et al. 2019), we found a substantial increase in LANTERN’s prediction accuracy for G_∞_ (Supplemental Fig S6). Overall, LANTERN provides the most accurate predictions of any single method across a broad benchmark of GPL datasets.

**Figure 3:**
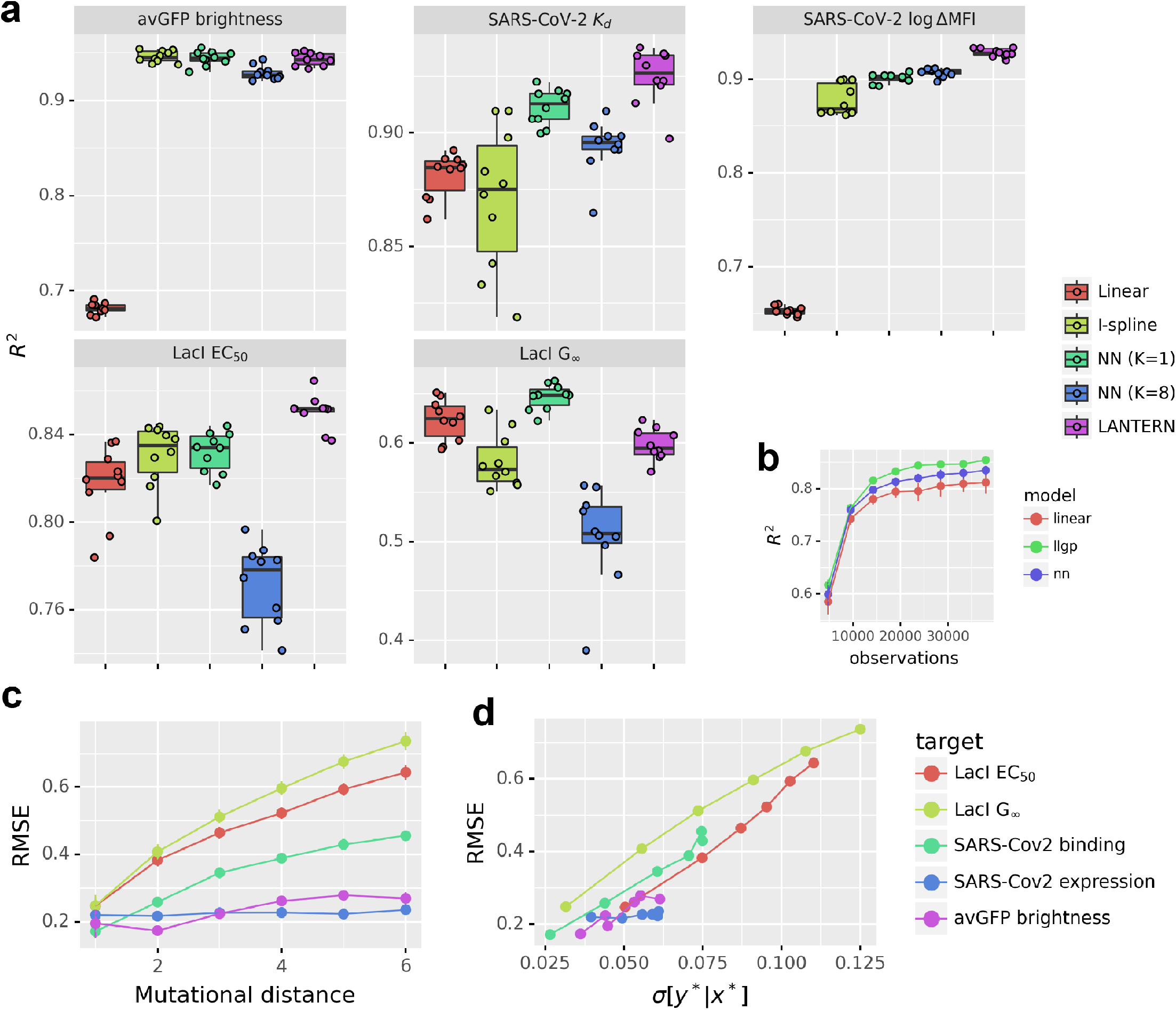
LANTERN equals or outperforms alternative models in predictive accuracy. (a) Ten-fold cross validated predictive accuracy, as determined by ***R***^2^ for different models across high-throughput GPL measurements. Box plots show the distribution of ***R***^2^ values while scatter points mark the value for individual folds. (b) Ten-fold CV predictive accuracy as a function of number of observations for LacI EC_50_. (c) LANTERN root mean squared error (RMSE) as a function of mutational distance from the wild-type sequence. (d) RMSE versus the average posterior predictive uncertainty of LANTERN (see section 4.5) for varying mutational distance from the wild-type.

To determine the impact of sample size on LANTERN performance, we evaluated cross-validated accuracy as a function of dataset size (Fig 3b). Due to the formulation as a Bayesian model, LANTERN straightforwardly balances the complexity of the model against the available GPL data. Consequently, the same training procedure for LANTERN applies regardless of dataset size. The relative gains of LANTERN over alternative approaches become most pronounced as the number of measurements increases. For smaller datasets, e.g. less than five thousand genotypes, LANTERN performs equally well as a simple linear model. Importantly, LANTERN does not underperform the linear model as dataset size decreases, reflecting the ability of LANTERN to straightforwardly scale *down* to relatively small datasets. LANTERN even scales down to small-scale measurements with only five to seven mutations (Supplemental Fig S7).

### 2.4 LANTERN quantifies predictive uncertainty

As a Bayesian modeling approach, LANTERN directly quantifies the uncertainty of its predictions. In order for these uncertainties to provide useful information, however, they must accurately reflect the degree of certainty that should be placed in each prediction when tested against real data (Snoek et al. 2019). We therefore compared the prediction uncertainties reported by LANTERN to its actual prediction error. The actual prediction error typically increase with increased mutational distance from the wild-type genotype (Fig 3c). This is due at least in part to the decrease in measurement coverage for variants with more mutations. A well-calibrated prediction uncertainty should similarly increase with mutational distance to reflect the decrease in available data. LANTERN’s predictions directly reflect this phenomenon, with uncertainty growing proportional to the prediction error (Fig 3d). LANTERN does appear to underestimate the overall true error rate however, a known issue with variational methods (Blei et al. 2016). Overall, the degree of confidence placed in each prediction by LANTERN is automatically specified by its uncertainty.

### 2.5 LANTERN provides interpretable models of genotype-phenotype landscapes

While LANTERN generates reliable predictions, it also straightforwardly explains these predictions through its interpretable components. Here, we demonstrate how this interpretability can be leveraged to understand the models learned by LANTERN for the three large-scale GPL datasets: avGFP, LacI, and SARS-CoV2. Across these datasets, the latent dimensionality learned by LANTERN ranged three to five (Fig 4a). We focus our analysis on the two most relevant dimensions of each dataset because additional dimensions capture an exponentially decreasing fraction of the total variation in the latent space of mutational effects, and correspondingly represent more minor details of each landscape (Supplemental Figs S8—S11).

**Figure 4:**
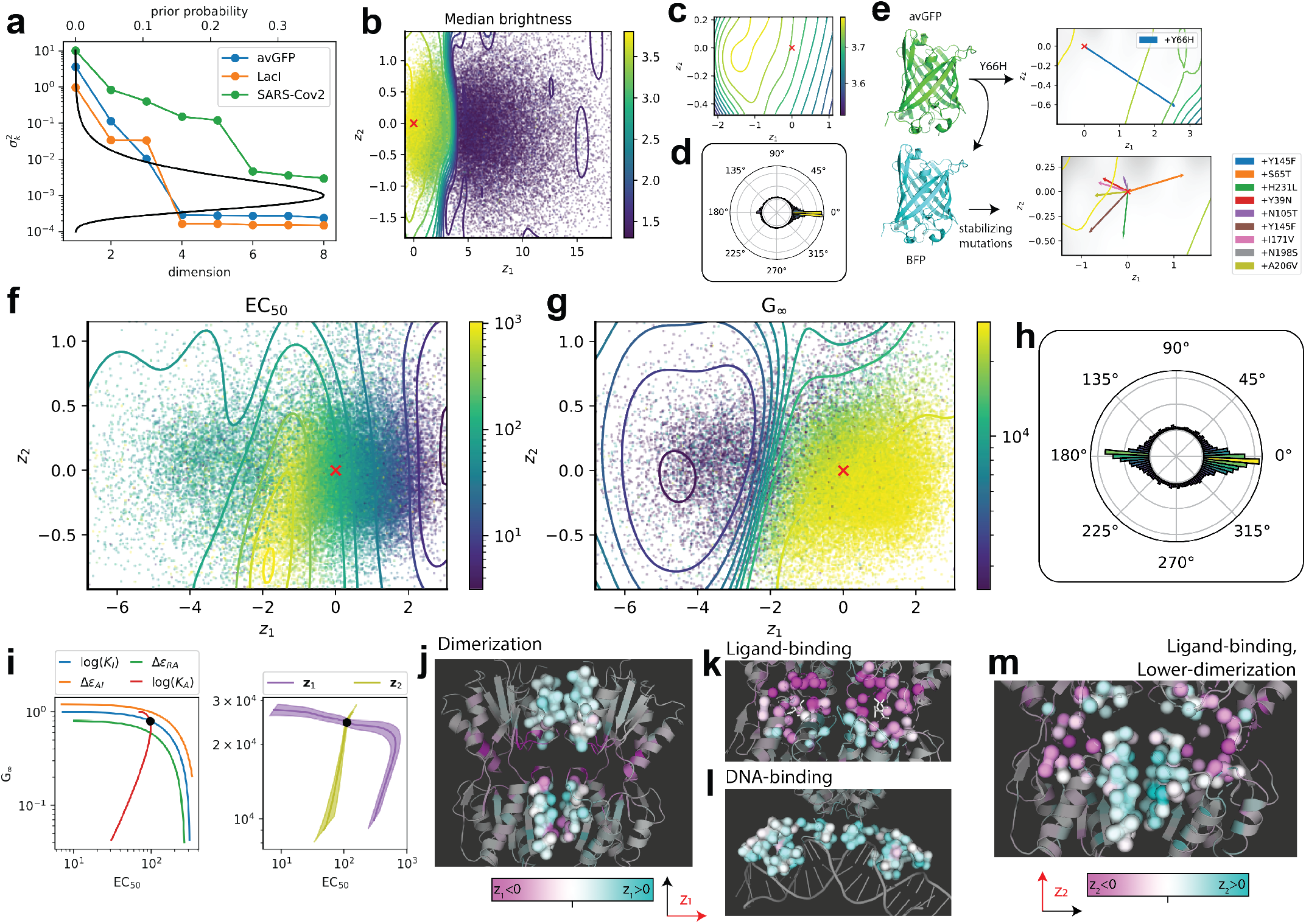
Interpretable models of avGFP brightness and LacI allostery. (a) Variance of each dimension in each large-scale GPL measurement. avGFP and LacI have three dimensions, while SARS-CoV-2 has five. (b) Learned avGFP brightness surface *f* (*z*) along the two highest relevance dimensions learned by LANTERN. Contours show equal values of the posterior mean of *f* (*z*), scatter points show the learned posterior mean of observations colored by their observed brightness, and red cross marks the wild-type origin (**z**_wt_ = **0**). (c) Focused region of the fluorescence surface, highlighting the region of maximal brightness off-center from the wild-type (red cross). (d) The distribution of latent, single mutant directions learned by LANTERN across the two highest relevance dimensions. (e) Analysis of mutations in blue fluorescent variants of avGFP. Blue fluorescent protein (BFP) is created by the Y66H mutation in avGFP. The latent mutational effect vector of Y66H is shown in the top panel. Contour lines show the posterior mean of *f* (*z*). Additional mutations in BFP that increase fluorescence are predicted to similarly increase avGFP fluorescence. These mutations may then have general structural stabilizing effects independent of fluorescent color. (f,g) The learned joint surface of LacI EC_50_ (f) and G_*∞*_(g). Contours show constant posterior mean of *f* (*z*), scatter points are the posterior mean *z* value of variants in the dataset colored by their observed parameter values, and red cross marks the wild-type origin. (h) The distribution of latent, single mutant effect directions across **z**_1_ and **z**_2_ learned by LANTERN for LacI. The majority of mutational effects (1374/2510 mutations within ∡ ≤ 25° of **z**_1_) lie near the positive (804 mutations) or negative (570) **z**_1_ axis. (i) Comparison of relationship between EC_50_ and G_*∞*_ when varying biophysical constants in an anlytic model of allostery (left) and over **z**_1_ and **z**_2_ (right). Each line represents the joint predicted value of EC_50_ and G_*∞*_ as the corresponding biophysical constant or latent dimensionsal effect (**z**_1_ or **z**_2_) is varied. Plots for **z**_1_ and **z**_2_ show predicted mean as solid line and 95% credible interval as a shaded region. (j—m) Association of latent mutational effects in **z**_1_ (j—l) and **z**_2_ (m) to different regions of the LacI protein. Highlighted regions are shown as connected surfaces in each panel. Residues are colored by their average posterior mean for the corresponding latent effect dimension. Highlighted regions are the dimerization interface (j), ligand binding pocket (k), DNA-binding domain (l), and a mixture of ligand-binding and lower-dimerization interface (m). All structures represent the active, DNA-bound state, with the exception of (k) which shows the inactive, ligand-bound state. In each panel, two LacI proteins are shown as a dimer.

#### avGFP

To determine how LANTERN makes predictions about avGFP fluorescent brightness, we analyzed the latent mutational effects and surface learned by the model. A sharp boundary along the most relevant dimension, **z**_1_, divides the latent mutational effect space into two regions: one with near wild-type fluorescent levels, and a second with complete loss of fluorescence (Fig 4b). Maximum brightness occurs in a region near (but not centered on) the wild-type (Fig 4c). Over 78% of mutations (1476/1879) decrease fluorescence because their mutational effect vectors point away from this maximum towards the region of decreased fluorescence (Fig 4d). Previous analysis of this landscape demonstrated that structural stability constituted a major factor in decreased fluorescence, with small increases in folding free energy leading to complete loss of functionality (Sarkisyan et al. 2016). However, discovery of this association depended on fixing the dimensionality of the neural network model used for the analysis. In contrast, LANTERN learns this association directly from data, finding a primary axis of decreased fluorescence along **z**_1_ that recapitulates the relationship between structural stability and brightness.

In order to further understand the LANTERN model of avGFP fluorescence, we analyzed mutations used in engineered variants of blue fluorescent protein (BFP, Yang et al. 1998). The foundational mutation of all BFP variants is Y66H. GFP variants containing Y66H have on-average fluorescence equal to 0.081 ± 0.010 the wild-type level and LANTERN coordinately predicts Y66H to decrease brightness, as expected in an assay to detect green fluorescence (Fig 4e). However, newer variants of BFP include additional mutations that increase the blue fluorescence, possibly through stabilizing protein structure (Ai et al. 2007). Interestingly, LANTERN predicts all of these mutations (with one exception: S65T) to similarly increase wild-type avGFP fluorescence. Additionally, these mutations point towards the maximum predicted brightness avGFP (Fig 4c). According to LANTERN then, these mutations may generally improve brightness independent of fluorescent color, possibly through improved structural stability.

#### LacI

Tack et al. (2021) measured the allosteric landscape of over 47,000 LacI variants with Hill-equation-like dose-response curves. With LANTERN, we modeled the landscape of these response curves with a multivariate phenotype of EC_50_ and G_∞_ for each variant. From the LANTERN model, the LacI GPL has a dimensionality of three, and the majority of mutational effects closely aligned with the positive or negative **z**_1_ axis (Fig 4a, f–h).

To interpret the latent mutational effect space learned by LANTERN, we compared the effects of changes in the two most relevant latent effect dimensions (**z**_1_ and **z**_2_) with the expectations from an analytic biophysical model for LacI dose-response (Razo-Mejia et al. 2018). Increases in **z**_1_ corresponds to a decreased EC_50_ and a slightly increased G_∞_. Conversely, decreases in **z**_1_ correspond to an increased EC_50_ and a sharply decreased G_∞_ (Fig 4f,g,i). This relationship between EC_50_ and G_∞_ along **z**_1_ is consistent with changes to three biophysical constants of the analytic model (which are indistinguishable with respect to their effects on EC_50_ and G_∞_, Fig 4i): the allosteric constant (Δ*∈*_AI_), the DNA operator affinity (Δ*∈*_RA_), or the ligand binding affinity of the inactive (non-operator binding) state (*K*_*I*_) (Eqs. 21, 22). So, we can interpret the changes in the **z**_1_ direction as free energy changes related to those three biophysical constants. Changes along the **z**_2_ dimension have a different effect: the positive **z**_2_ direction corresponds to a decreased G_∞_ at roughly constant EC_50_ (Fig 4f,g,i), and the negative z2 direction corresponds to EC_50_ and G_∞_ that remain near the wild-type values. This is consistent with changes to a fourth biophysical constant of the analytic model: the ligand binding affinity of the active (operator binding) state (*K*_*A*_, Fig 4i). So, we can interpret the changes in the **z**_2_ direction as free energy changes related to the active state ligand affinity.

To further understand the connection between the latent mutational effects and the biophysics of LacI, we analyzed the distribution of mutational effects in the **z**_1_ and **z**_2_ dimensions across the LacI protein structure. Mutations in the dimerization region of LacI generally increase **z**_1_ (Fig 4j). Since mutations in this region are unlikely to affect the DNA operator affinity or ligand affinity, these shifts in **z**_1_ could be due to changes in the allosteric constant (Δ*∈*_AI_). Conversely, mutations in the ligand-binding region of LacI more often decrease **z**_1_ (Fig 4k). Mutations in this region might be expected to affect the ligand affinity (*K*_*I*_), though previous work indicates that mutations near the ligand-binding region can also affect the allosteric constant (Chure et al. 2019). Finally, mutations in the DNA-binding domain of LacI generally increase **z**_1_ (Fig 4l). This is consistent with a decrease in the DNA operator affinity (Δ*∈*_RA_). Surprisingly, mutations that increase **z**_2_, corresponding to an apparent decrease in the active-state ligand binding constant (*K*_*A*_), are not typically found near the ligand-binding region. Instead, they largely occur at the N-terminal half of the dimerization interface (Fig 4m). This region of the protein undergoes a conformational shift when the protein switches between the active and inactive states (Lewis et al. 1996). So, mutations here could affect *K*_*A*_ via changes in allosteric communication between the DNA-binding and ligand-binding domains of the protein.

#### SARS-CoV-2 Receptor Binding Domain

Starr, Greaney, et al. 2020 measured the effect of over 3,800 distinct mutations in the SARS-CoV-2 spike protein receptor binding domain (RBD) for their binding affinity to ACE2 (log *K*_*d*_) and structural stability (measured as change in yeast display expression, Δ log MFI), sampling in total more than 170,000 unique variants. LANTERN discovered five dimensions in this landscape measurement (Fig 4a). The most common direction of mutational effects roughly follows a gradient of steepest descent for structural stability measured by Δ log MFI (Fig 5a–c). We derived an axis from this direction, which we refer to as the *stability* axis (Fig 5c). The direction of latent effect for nearly 50% of mutations (1842/3798) lie within ten degrees of this axis. We next identified a nearly orthogonal axis that lies along a constant ridge of RBD structural stability, which we call the *binding* axis (Fig 5c). ACE2 binding decreases along both axes, but structural stability only changes along the *stability* axis (Fig 5d—f). This suggests that mutations along the *stability* axis, i.e. most mutations of the RBD, decrease structural stability. This decrease in structural stability then disrupts RBD binding to the ACE2 receptor, since the spike protein must fold correctly before it can bind to ACE2. Conversely, mutations along the *binding* axis do not impact structural stability and may be particularly important in forming the RBD-ACE2 complex independent of structural stability. This interpretation is supported by the location of mutations within the different protein structure domains: the majority of mutations with latent effects along the *binding* axis are near the RBD-ACE2 interface while mutations along the *stability* axis are distributed throughout the core RBD domain (Fig 5f,g).

**Figure 5:**
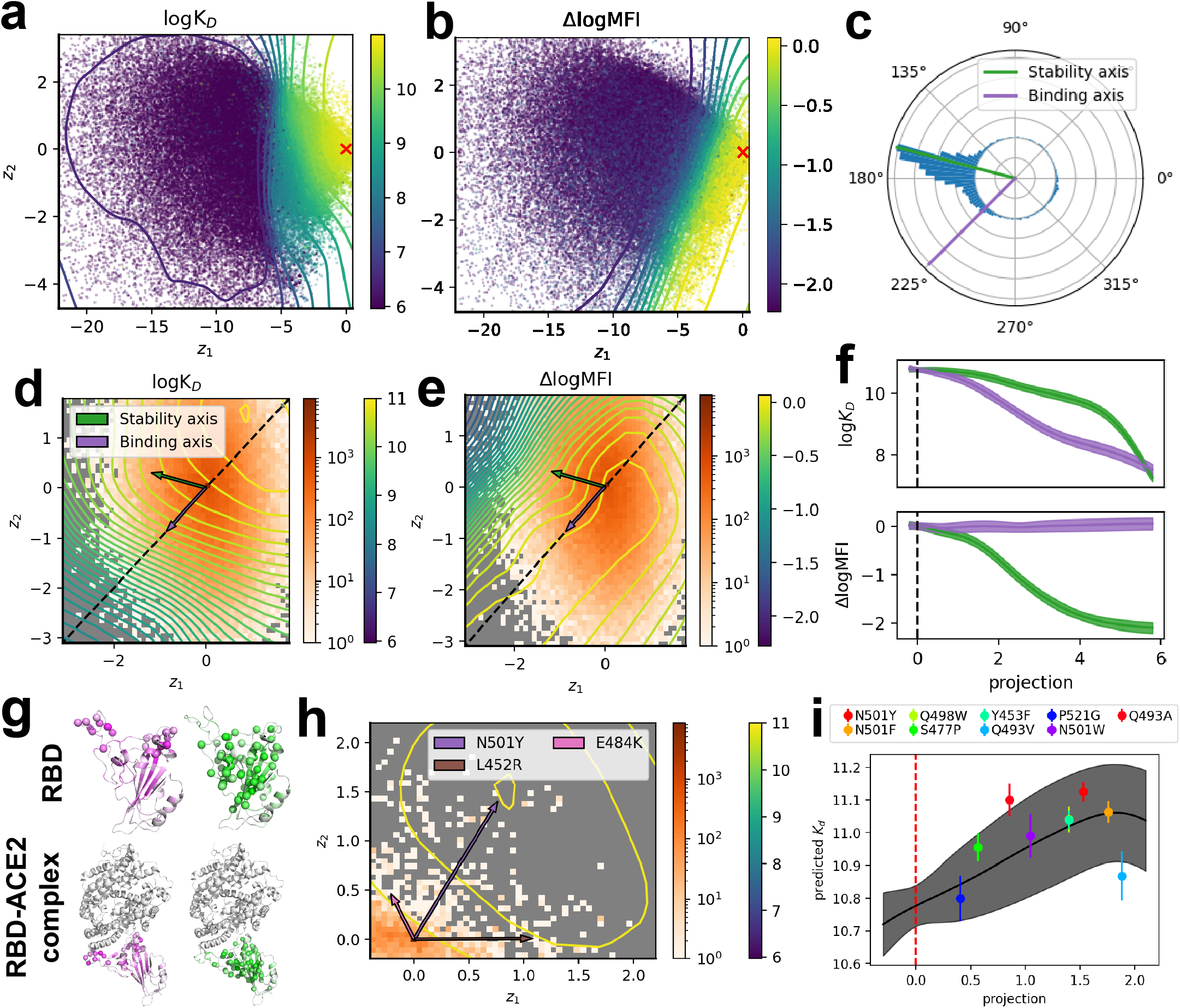
SARS-Cov2 joint binding and expression landscape. (a, b) Learned surface *f* (*z*) for SARS-Cov2 binding (a) and expression (b). Contours show posterior mean of *f* (*z*), scatter points are mean posterior *z* for individual variants colored by their observed mean value, and red crosses mark the wild-type origin. (c) Distribution of latent, single mutant effect directions along the first two dimensions of **z**, with *stability* axis (green) and *binding* axis (purple). (d, e) Joint binding and expression surfaces with identified axes overlayed. Contours are the mean of the variational posterior of *f* (*z*) and the two-dimensional histograms mark the density of observed variants in *z*-space. (f) The predicted binding and expression as a function of the projection along both identified axes. (g) Structural association of the identified axes. SARS-Cov2 binding the ACE2 RBD and the SARS-Cov2 protein in isolation are shown. Colors indicate the average value of mutational effect projections onto the corresponding axis. Spheres indicate a strong average alignment (∡≤ 36°) between mutational effects at a residue and the corresponding axis. (h) Mutations associated with COVID-19 outbreaks (Hodcroft 2021). Individual mutational effects are shown as vectors. Contours and histogram are as in (d,e). (i) Predicted binding along the mutational effect direction of N501Y. Projection corresponds to the scale factor of the N501Y unit direction vector, solid black line is the posterior mean of *f* (**z**), shaded regions is the 95% confidence interval, and solid red line is the wild-type origin. Single-mutant variants that align with the virulence axis and have a positive projection value are shown with error bars representing a 95% confidence interval of the single mutant measurement.

To demonstrate what LANTERN can reveal about clinically relevant mutations, we analyzed mutations that have been found in recently identified SARS-CoV-2 variants of concern (Hodcroft 2021; Harvey et al. 2021, Fig 5h). The latent mutational effects of these mutations all point in distinct directions within the latent space learned by LANTERN, and each direction corresponds to higher binding affinity than the wild type. So, although each of these mutations has a distinct impact on the protein’s function in terms of latent mutational effect, each is predicted to increase ACE2 binding affinity. One mutation in particular, N501Y, has a mutational effect vector that points directly toward the predicted maximum of RBD-ACE2 binding affinity. N501Y occurs in the B.1.1.7 variant that has steadily increased in proportion worldwide (Centers for Disease Control and Prevention 2021; Frampton et al. 2021). Given the importance of this mutation to the ongoing pandemic, we analyzed the ten mutations with the most similar latent effects, determined by the similarity of their latent mutational effect directions to that of N501Y. Among these mutations, RBD-ACE2 binding strength increases, and measurements generally confirm stronger ACE2 binding for single-mutants with these mutations (Fig 5i). The LANTERN analysis indicates that these mutations involve similar mechanisms as N501Y, based on the similarity their latent mutational effects. So, these mutations may be particularly important for genomic surveillance of SARS-CoV-2.

### 2.6 LANTERN quantifies local robustness and additivity

Despite the complexity of GPLs, quantitative metrics can summarize their most important features and simplify their analysis. For example, global GPL metrics provide insight into adaptive evolution and engineering potential (Kondrashov et al. 2015). With LANTERN, we define two novel metrics based on local properties of the landscape: slope and curvature. The slope of the landscape describes the rate of change of the surface at each position in latent mutational effect space (Fig 6a). With zero slope, the phenotype remains constant in response to small latent mutational effects. So, the (inverse) slope is associated with *robustness*. The curvature of the landscape reflects the rate of change of the slope, and zero curvature implies that mutations have a constant effect on phenotype (Fig 6b). In regions with zero curvature, mutations have no epistasis. So, the (inverse) curvature is associated with *additivity*.

**Figure 6:**
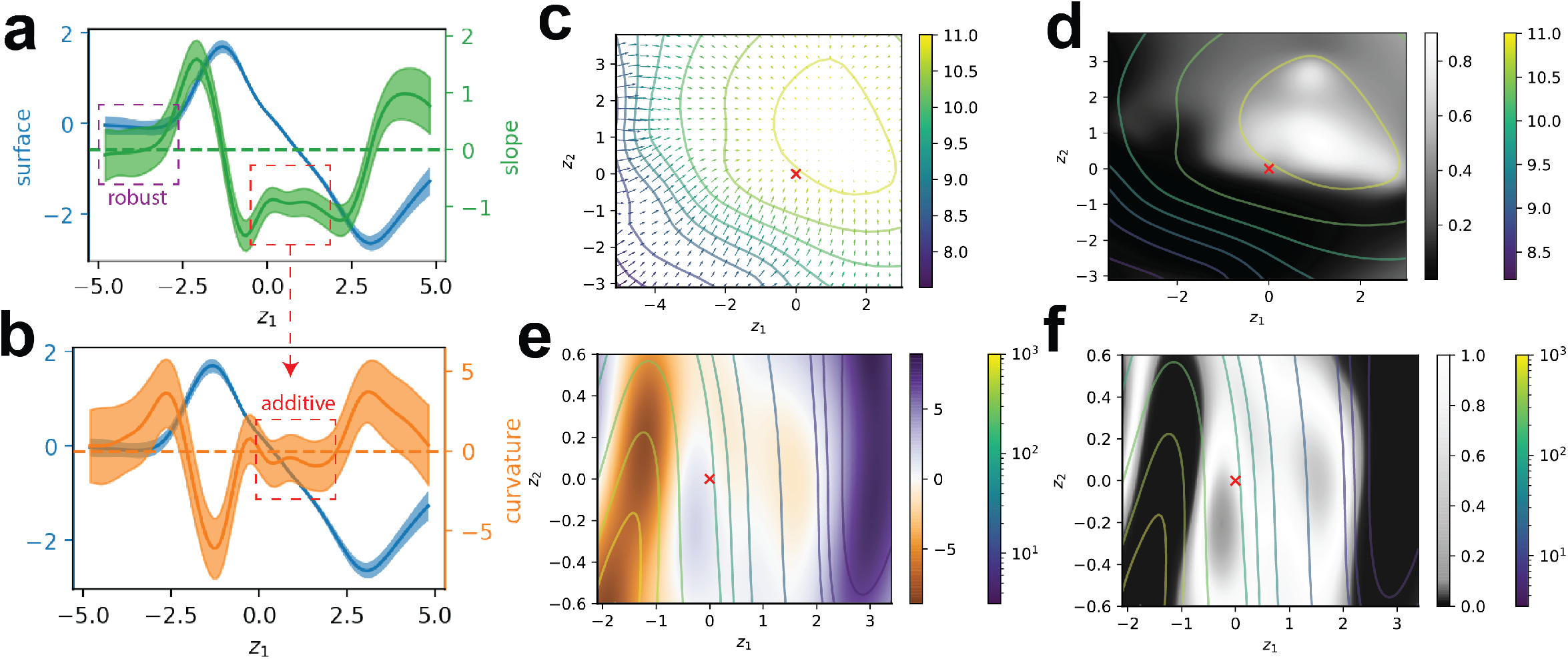
Local *robustness* and *additivity* of GPLs. (a) LacI EC_50_ surface and slope along **z**_1_. Posterior of *f* (**z**) is shown in blue and slope 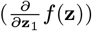 is shown in green. When the slope is zero, the surface is locally *robust* (purple box). (b) LacI EC_50_ surface and curvature along **z**_1_. Posterior of *f* (**z**) is shown in blue and curvature 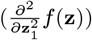 is shown in orange. When the curvature is zero, the surface is locally *additive* (red box). A curvature of zero also implies the slope is constant, though may be non-zero (a, red box). In both (a) and (b), solid lines are posterior mean and shaded regions are 95% credible intervals of *f* (**z**). (c) The gradient of SARS-CoV-2 binding. Arrows show the posterior mean of the gadient (∇ *f* (**z**)) and the contours mark the posterior mean of ∇*f* (**z**). The multidimensional equivalent of *robustness* is when *f* (**z**) = 0. (d) The *robustness* of SARS-CoV-2 binding. Values near one indicate near zero gradient. (e) The curvature of LacI EC_50_ in multiple dimensions, calculated as the Laplacian (Δ*f* (**z**), see methods). The multidimensional equivalent of *additivity* is when Δ*f* (**z**) = 0. (f) The *additivity* of LacI EC_50_. Values near one indicate Δ*f* (**z**) is close to zero.

To apply the concept of robustness to the multidimensional mutational effect spaces learned by LANTERN, we generalized the slope to multiple dimensions as the surface gradient ∇*f* (**z**). The gradient represents the rate of change in each dimension as a vector and a gradient with values near zero represents multidimensional robustness (Fig 6c). In the case of SARS-CoV-2, LANTERN predicts high robustness near the predicted maximum of binding strength between the RBD and ACE2 (Fig 6d). This region of latent mutational effect space poses a potential clinical threat, as it represents genetically stable SARS-CoV-2 infectivity through strong affinity between the RBD and ACE2. Variants in this region may then have more mutational flexibility to evade immune response while maintaining infectivity (Hie et al. 2021; Atlani-Duault et al. 2021; Harvey et al. 2021).

To similarly quantify additivity in multiple dimensions, we add the curvature across each dimension to compute the Laplacian (Δ*f* (**z**), Fig 6e). A value of zero for the Laplacian indicates a constant rate of change in all dimensions and constitutes multidimensional additivity. As an evaluation of additivity as a useful metric, we analyzed the additivity surface of LacI EC_50_ (Fig 6f). Previous analysis of the LacI EC_50_ phenotype showed that mutational effects in single and double mutants combine with no epistasis (Tack et al. 2021). The additivity surface quantitatively represents this phenomenon, with the EC_50_ surface having high additivity around the wild-type (Fig 6f). Notably, we draw this conclusion directly from the additivity surface predicted by LANTERN, rather than through combinatorial screening of epistatic effects in single and double mutants.

## 3 Discussion

LANTERN addresses the need for GPL models that make accurate predictions while remaining highly interpretable. We show that in a benchmark across multiple GPLs, LANTERN achieves equal or better predictive accuracy compared with state-of-the-art DNN models (Fig 3). However, our comparison to DNN models may remain incomplete because changes in network architecture or training hyperparameters can marginally improve predictive accuracy (Bergstra et al. 2012; Balaprakash et al. 2019; Kandasamy et al. 2018). This further highlights the advantages of LANTERN though, because LANTERN provides a single modeling interface for *any* GPL measurement with no additional tuning necessary. Additionally, DNNs typically require large-scale measurements to ensure satisfactory performance but provide no clear cutoff for how much data is sufficient. In contrast, LANTERN scales to any dataset size (Supplemental Fig S7). Overall, we expect LANTERN will provide accurate predictions in a broad class of GPLs and impact engineering and research endeavors that depend on extrapolation to new genotypes.

Typically, improvements to model predictive accuracy come at the expense of model interpretability, with an assumption that this trade-off is unavoidable (Lipton 2018). In the case of GPLs, LANTERN shows that this trade-off is unnecessary: optimal predictions do not preclude interpretable modeling. Specifcally, due to its construction from easily understandable components, LANTERN automatically provides an explanation for every prediction it makes. Furthermore, we establish that these components also independently provide insight into the biological mechanisms of GPLs. For example, LANTERN learns latent mutational effect spaces that reflect the underlying biophysical process, possibly up to a rotation (Fig 2e). LANTERN also presents straightforward explanations of the effects of individual mutations through their latent effect vector. These vectors facilitate clear understanding of how each mutation contributes to an observed phenotype (Fig 4e, 5h) and how these mutations are tied to structural biochemisty (Fig 4j—m, 5g). Finally, LANTERN quantifies the new metrics of local *robustness* and *additivity* that provide novel perspectives for understanding large-scale GPL measurements (Fig 6). Overall, we expect LANTERN’s interpretability will enhance our understanding of diverse landscapes across biology.

LANTERN models GPLs as a non-linear surface over a low-dimensional latent mutational effect space, a form of epistasis where non-additivity arises from global structure (Starr and Thornton 2016; Jakub Otwinowski et al. 2018). Certain theoretical models of GPLs, which until recently could not be verified due to the lack of sufficient experimental data, similarly concentrated on the existence of low-dimensional manifolds that explain the complexity of GPLs (Orr 2005; Tenaillon 2014). These models suggest that GPLs will commonly involve low-dimensional structure due to the benefits in adaptive evolution (Sato et al. 2020; Husain et al. 2020). LANTERN will therefore likely have broad applicability across GPL measurements of diverse biological systems (Kinney et al. 2019). Additionally, LANTERN extends beyond existing global epistasis modeling approaches that suffer from restrictive assumptions, e.g., a predetermined dimensionality or a fixed family of non-linear functions (Jakub Otwinowski et al. 2018; Sailer, Shafik, et al. 2020) (Supplemental Figs S12, S13). LANTERN instead learns these details directly from the data. This enables direct comparisons between GPLs that are not possible with other modeling approaches, like the relative complexity of two landscapes quantified by their dimensionality (Fig 2b, 4a).

Large-scale GPL measurements will increasingly influence bioscience initiatives of the future. Despite the growing magnitude of experimental throughput, the full genotypic space will always remain under-sampled. To overcome this fundamental limitation, LANTERN facilitates progress through reduction of landscapes to their minimal complexity. In this way, LANTERN transforms the intractable challenge of exhaustive genotypic sampling to a manageable exploration of a low-dimensional space. GPL-enabled investigations can then explore this space in an efficient manner, relying on LANTERN’s guidance toward uncharted regions of phenotypic diversity.

## 4 Methods

### 4.1 Genotype-phenotype landscape datasets

We aggregated GPL measurements from published sources (Sarkisyan et al. 2016; Tack et al. 2021; Starr, Greaney, et al. 2020). In the case of LacI, we filtered observations to variants with Hill-equation-like dose-response curves to avoid inference on inaccurate hill parameter estimates. We combined the two libraries of SARS-Cov-2 for each of binding and expression measurements to make a single, aggregate dataset for both measurements. In the case of joint modeling, we merged the binding and expression datasets for cases where a variant was present in both.

We prepared all training datasets in a similar fashion. Within each dataset, we one-hot encoded all mutations (Angermueller et al. 2016). Specifically, for *p* total mutations in a dataset (Table 1), each variant *i* was represented as the one-hot encoded vector *x*_*i*_ ∈ [0, 1]^*p*^. For each corresponding phenotype *y*_*i*_ ∈ ℝ^*D*^ with phenotype dimensionality *D*, we standardized each phenotype dimension seperately to a mean of zero and standard deviation of one. The final dataset for training was then 𝒟 = {*x*_*i*_, *y*_*i*_}|1 ≤ *i* ≤ *N*] for *N* total variants.

### 4.2 Hierarchical Bayesian modeling of GPLs

We constructed LANTERN with two key components: a latent surface *f* (**z**) and a set of latent mutational effects, represented by a matrix *W* = [*z*^(1)^, *z*^(2)^, …, *z*^(*p*^)] ∈ ℝ^*K*×*p*^ where *z*^(*i*)^ represents the mutational effect vector of mutation *i* (Eq 3, Fig 1a). First, we placed a Gaussian process (GP) prior on the latent surface *f* (**z**):

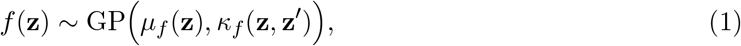

with mean and kernel functions *µ*_*f*_ and *κ*_*f*_, respectively. We set the mean function *µ*_*f*_ as an unknown constant value, 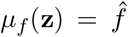. The kernel function *κ*_*f*_ describes the covariance between different observations of *f* : cov *f* (**z**), *f* (**z**^**′**^) = *κ*_*f*_ (**z, z**^**′**^)). We used the rational quadratic covariance function:

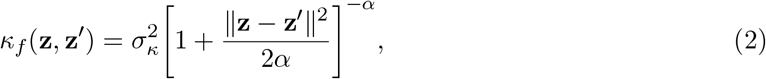

where *σ*_*κ*_ is an unknown scale parameter and *α* controls the overal distribution between smooth and rugged regions of *f*. We did not include a lengthscale parameter for the norm ∥**z** − **z**^**′**^∥ used in the kernel because we already learn the relative scale between dimensions through the hierarchical prior on dimension variance (Snelson et al. 2006).

Next, we specified the hierarchical prior for the unknown mutation effects *W* ∈ ℝ^*K*×*p*^ for *K* latent mutational effect dimensions and *p* mutations. For a variant *i* with mutation vector *x*_*i*_ ∈ {0, 1}^*p*^ (see Sec 4.1), the latent mutational effect vector *z*_*i*_ (conditional on *W*) is computed as

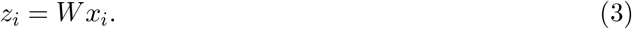

Note that *z*_*i*_, representing the combination of latent mutational effects for each mutation in variant *i*, is distinct from a dimension of the latent mutational effect space (e.g. **z**_1_).

For each row *r* of *W, w*_*r*_, we defined a Gaussian prior for each element of *w*_*r*_:

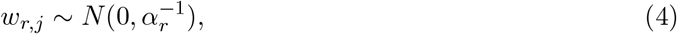

where *α*_*r*_ is the precision (or inverse variance) of dimension *i*. We assumed each *α*_*r*_ to be drawn from a Gamma distribution (Bishop 1999):

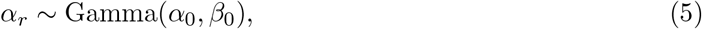

where *α*_0_ and *β*_0_ are model hyper-parameters. We set *α*_0_ = *β*_0_ = 10^−3^ in all experiments here, which generally leads to models that minimize the number of effective dimensions (Bishop 1999). Different values of *α*_0_ and *β*_0_ would change the scale learned for the latent dimensional effect space (Supplemental Fig S14). However, the overall scale of the latent dimension is not relevant to the model. Instead, a certain small number of dimensions should have substantially larger magnitude than the rest. So, we use the prior for *α*_*r*_ to ensure that most dimensions are effectively removed from the model while allowing a small number of dimension with non-negligible variance (1*/α*_*r*_). We rank the importance of each dimension by its variance, which we refer to as its *relevance*.

To combined these components when modeling a GPL dataset 𝒟, we assume that each phenotype *y*_*i*_ is conditionally independent given the unknown variables:

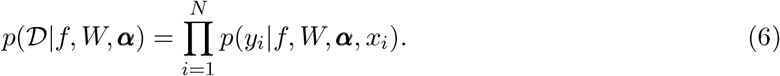

The likelihood of each phenotype *y*_*i*_ was calculated as

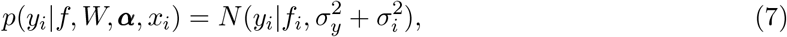

where 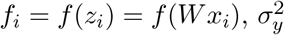 is an unknown variance parameter estimated from the data and 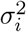 is any additional measurement uncertainty provided for each variant in the dataset.

We treat the kernel parameters *σ*_*κ*_ and *α*, as well as the global phenotype noise, 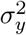 as unknown variational parameters (Section 4.3) constrained to be positive. These parameters are then still learned for each dataset.

### 4.3 Approximate inference

#### Variational inference

Given the specified model, inference involves the recovery of the posterior distribution:

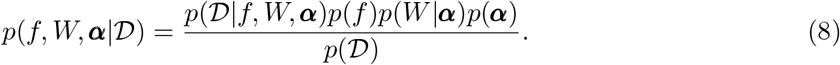

The exact posterior is analytically intractable, and we instead relied on an approximation through variational inference (VI) (Blei et al. 2016). VI recasts the inference procedure as an optimization problem to minimize the KL divergence between an approximate posterior *q*(*f, W*, ***α***) and the true posterior *p*(*f, W*, ***α*** |𝒟). We take the typical mean field approach which assumes that *q* decomposes multiplicatively across unknown variables: *q*(*f, W*, ***α***) = *q*(*f*) ∏_*i*_ [*q*(*α*_*i*_) ∏_*j*_ *q*(*w*_*ij*_)]. See below for the choice of *q* for each variable in the model. Returning to the objective, which is still intractable, we instead minimized it indirectly through maximization of the evidence lower bound (ELBO, Supplemental Fig S16):

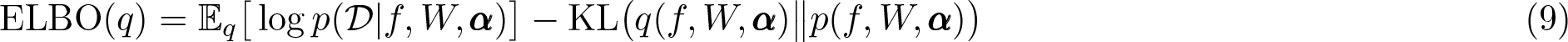

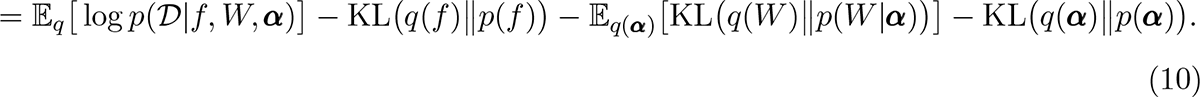

This objective bounds the model evidence *p*(𝒟) up to an unknown constant and therefore maximizing the ELBO corresponds to minimizing the KL divergence between *q* and the true posterior.

In order to make inference scalable to large-scale GPL datasets, we employed stochastic variational inference (SVI) (Hoffman et al. 2013). SVI approximates exact variational inference by sub-sampling the training data in smaller mini-batches and computing a noisy approximation of the gradient of the ELBO:

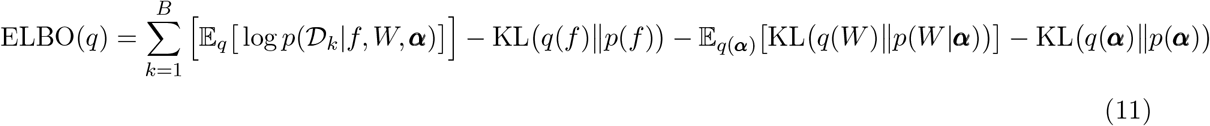

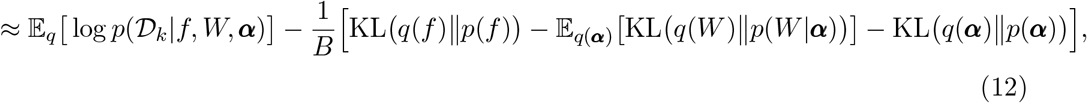

where 𝒟_*k*_ is the *k*th mini-batch and there are *B* total mini-batches.

We rely on variational distributions *q* for which the KL divergence is analytically tractable (see below). But, computation of the expected log-likelihood of the data remains an analytical challenge. So, we approximate the true expectation through unbiased Monte-Carlo samples. This is made possible by re-parameterizing each variational distribution *q*(*θ*) in terms of a new distribution *q*(*∈*) such that *θ* = *g*(*∈*) and 𝔼_*q*_[*h*(*θ*)] = 𝔼_*∈*_[*h*(*g*(*∈*))]. We can then approximate the true expected log-likelihood as

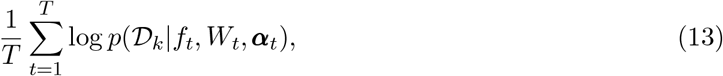

for *T* Monte-carlo samples (we set *T* = 1 in all cases) and Monte-carlo samples of each parameter using its reparameterized distribution: *θ*_*t*_ = *g*(*∈*_*t*_) and *∈*_*t*_ ∼ *q*(*∈*). This greatly reduces the variance in gradient calculations and improves inference (Diederik P Kingma et al. 2013; Blundell et al. 2015).

#### Variational distributions

We used a variational distribution of *f* (**z**) using the methods of (Hensman et al. 2015). Briefly, the complete distribution on *f* is approximated by learning the distribution of *f* at specific inducing points *Z* (which are parameters of the variational distribution and not random variables) and unknown function values *u* = *f* (*Z*). A variational distribution on *u* is defined as *q*(*u*) ∼ *N* (*µ*_*u*_, Σ_*u*_), the variational distribtuion on *f* is

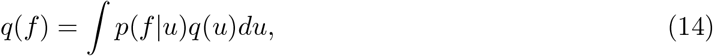

and can be analytically calculated from the GP prior. For *M* inducing points, the computational cost of this approximation during inference is *O*(*M* ^3^), realizing substantial savings when *M* ⋘ *N*. We use the implementation of the approximation approach for *f* (**z**) provided by the python library `gpytorch` (Gardner et al. 2018).

For the remaining parameters, weights *w*_*i,j*_ and precisions *α*_*i*_, we set their variational family to the same family as their corresponding prior:

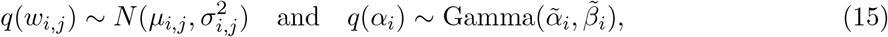

where 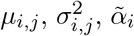, and 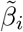 are optimized with respect to the ELBO (Eq. 12). Note that *p*(***w***_*i*_|*α*_*i*_) = *N* (***w***_*i*_|0, *α*_*i*_) depends on *α*_*i*_, which we obtain from a Monte Carlo sample from *q*(*α*_*i*_).

We implemented LANTERN with the automatic differentiation library `pytorch` (Paszke et al. 2019), with GP components of the model relying on `gpytorch` (Gardner et al. 2018). We trained models on compute systems with NVIDIA TITAN V GPUs, with ≥ 12 GB of memory and 5120 CUDA cores. We trained LANTERN models with mini-batch size 4096 or 8192, depending on GPU capacity, learning rate of 10^−2^ using the Adam optimizer (Diederik P. Kingma et al. 2014), and for 5,000 epochs. We set the maximum number of latent mutational effects dimensions to eight. A total of 800 inducing points were learned for each model, with initial positions uniform randomly sampled over the range [−10, 10].

### 4.4 Computing *additivity* and *robustness*

*Additivity* and *robustness* were derived from the Laplacian (Δ*f* (*z*)) and gradient (∇*f* (*z*)) posterior predictive distributions, respectively (Section S3). For a location *z*_*k*_ in the latent mutational effect space, we calculate the analytic posterior distribution of the gradient, *q*(∇*f* (*z*_*k*_)) ∼ *N* (*µ*_∇_, σ_∇_), and Laplacian, 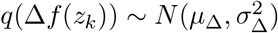 (See Equations S2 and S13, respectively). Both *robustness* and *additivity* were quantified from the unnormalized density, or kernel, of their respective differntial operator’s posterior predictive distribution at zero:

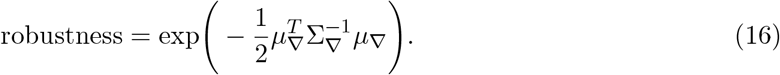

and

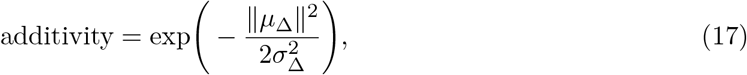

Values of *additivity* and *robustness* close to 1 then imply high probability of zero values for the underlying differential operators Δ*f* (*z*) and ∇*f* (*z*).

### 4.5 Comparison of predictive accuracy

#### Models

We employed the montonic I-spline model of (Jakub Otwinowski et al. 2018) via the `dms_variants` library (github.com/jbloomlab/dms_variants), which provides an implementation of the model in python. We trained monotonic spline models on each dataset using default parameters. We used a Gaussian likelihood for all datasets.

We adapted deep neural network architectures from recent publications performing regression on GPL measurements (Sarkisyan et al. 2016; Pokusaeva et al. 2019). These architectures follow a feed-forward structure:

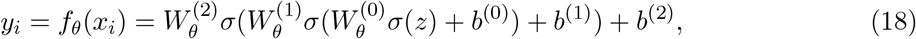

with weights *W* ^(*k*)^ and biases *b*^(*k*)^ and non-linearity *σ*. These models generally have a low-dimensional initial hidden layer, e.g. *W* ^(0)^ ∈ ***R***^*L*×*K*^ and *b*^(0)^ ∈ ***R***^*L*^ with *L* ⋘ *K*. The primary hidden layer has width *w*: *W* ^(1)^ ∈ ***R***^*w*×*L*^ and *b*^(0)^ ∈ ***R***^*w*^. A final linear layer transforms the hidden neurons to the output dimension: *W* ^(2)^ ∈ ***R***^*D*×*w*^ and *b*^(2)^ ∈ ***R***^*D*^. For this study, models were constructed with an initial latent dimensionality *L* of either one or eight, sigmoid nonlinearity *σ*, and hidden width of *w* = 32. We trained neural networks with the Adam optimizer, with mini-batch size of 128, learning rate of 10^−3^, for 100 epochs. We chose the epoch length as it minimized the held-out validation prediction error. We minimized the mean squared error, with losses weighted inversely proportional to measurement uncertainty.

#### Cross validation

We evaluated all models with ten-fold cross validation. The same folds for each dataset were re-used across modeling approaches in order to ensure accurate comparisons. We quantified predictive accuracy across folds with the coefficient of determination, 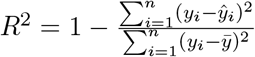 for model predictions *ŷ*_*i*_ and observations mean 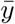. In the case of measurement uncertainty, the individual prediction scores were weighted proportionally to the certainty of their measurement:

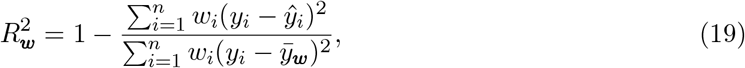

for weights 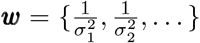 and 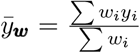. In the case of models that are approximated with variational methods, we made predictions with the approximate posterior mean of all unknown random variables.

In order to determine the relationship between predictive accuracy and dataset size, we performed cross-validation training with randomized subsets of the training data. We trained models from data subsampled at 5,000 increments of the full training set size, and evaluted performance on the held out test data for all resulting models.

#### Uncertainty quantification

We quantified predictive uncertainty of LANTERN with Montecarlo draws from the approximate variational posterior. Specifically, a Monte-carlo sample of the unknown mutation effects were taken from the variational posterior: 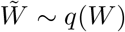 This sample was then used to calculated the latent position of every variant: 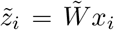. Then, the approximate posterior predictive distribution of *f* was used to sample a predictive phenotype for the variant: 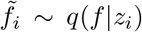. This two stage sampling process was repeated fifty times to estimate the overall uncertainty of *f*_*i*_ for the variant.

### 4.6 Simulated biophysical allosteric model

Simulated data was generated using a biophysical model for allosteric transcriptional regulation (Razo-Mejia et al. 2018). That model relates the dose-response curve, *f* (*c*), of an allosteric transcription factor to the biophysical parameters such as binding constants and free-energy differences between protein states:

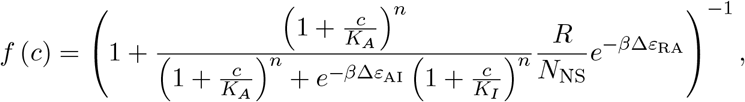

where *c* is the ligand concentration, *K*_*A*_ and *K*_*I*_ are the dissociation constants between the ligand and the transcription factor in the active and inactive states, respectively, Δ*ε*_AI_ is the free energy difference between the active and inactive states, Δ*ε*_RA_ is the free energy difference between specific and non-specific binding of the transcription factor to DNA, *R* is the number of transcription factor molecules in the cell, *N*_NS_ is the number of non-specific DNA binding sites, *β* = (*k*_*B*_*T*)^−1^, *k*_*B*_ is the Boltzmann constant, and *T* is the temperature. Simulated results were generated for the baseline dose-response, G_0_ = *f* (0), the saturated dose-response, G_∞_ = lim_*c*⟶∞_ *f* (*c*), and the sensitivity, EC_50_:

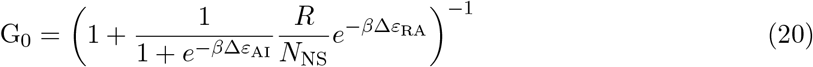

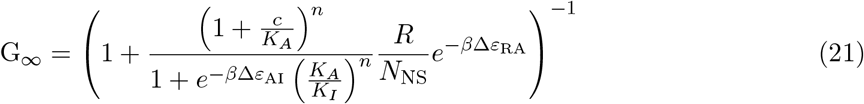

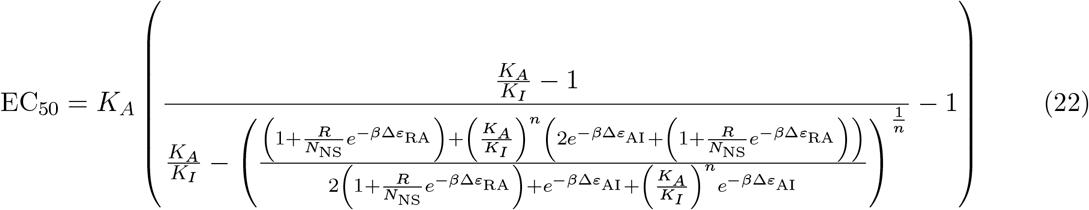

The values assumed for the wild-type parameters were: *K*_*A*_ = 139 *µ*mol/L, *K*_*I*_ = 0.53 *µ*mol/L, *β*Δ*ε*_AI_ = 4.5, ***R*** = 150, *n* = 2, *β*Δ*ε*_RA_ = −15.3, *N*_NS_ = 4.6 × 10^6^.

Simulated datasets were generated assuming a protein with 300 amino acid positions and six possible amino acid substitutions at each position.

Three different simulated datasets were generated:

1. 𝒟_1_: data for a 1D manifold, varying only the *β*Δ*ε*_AI_ parameter
2. 𝒟_2_: data for a 2D manifold, varying *β*Δ*ε*_AI_ and *β*Δ*ε*_RA_
3. 𝒟_3_: data for a 3D manifold, varying *β*Δ*ε*_AI_, *β*Δ*ε*_RA_, and log_10_ (*K*_*A*_)

For each dataset a 300 amino-acid length protein was simulated with 6 possible substitutions for each amino acid position. Random mutational effects were assigned to each possible substitution using the following procedure: First, a mean effect was chosen for each position as a random draw from a normal distribution with zero mean and standard deviation of 1.5. Then, the effect of each specific substitution was determined as the mean effect at the corresponding position multiplied by a random integer between -1 and 1 plus a random draw from a normal distribution with zero mean and standard deviation of 1.

For each simulated dataset, 100,000 simulated variants were chosen by first assigning the number of amino acid substitutions for each variant based on random draws from the empirical distribution from an experimental dataset (Tack et al. 2021). The positions and identities of simulated substitutions were then randomly chosen as unbiased random draws from the set of positions (without replacement), and possible substitutions (with replacement). The shifts of the biophysical parameters (*β*Δ*ε*_AI_, *β*Δ*ε*_RA_, and log_10_ (*K*_*A*_)) were then determined for each simulated variant by summing the effects of each substitution. The effects of changes to log_10_ (*K*_*A*_) were much stronger than the effects of the other parameters, so the shift in log_10_ (*K*_*A*_) for each variant in the 3^rd^ dataset was scaled by a factor of 0.1. The biophysical parameters for each variant were determined by adding the shift for that parameter-variant combination to the wild-type parameter values listed above. Finally, equations 20—22 were used to calculate values for G_0_, G_∞_, and EC_50_ for each simulated variant.

### Code availability

Source code is available at github.com/usnistgov/lantern.

## 5 Acknowledgements

We would like to thank Blaza Toman, Swarnavo Sarkar, and Dennis Leber for thoughtful feedback on this manuscript. PDT and ADP were supported through NRC Fellowships.

Analysis performed in part on the NIST Nisaba HPC clusters.

## 6 Author Contributions

PDT and DJR conceived of the approach

PDT implemented LANTERN and ran analyses DJR generated simulated biophysical landscapes PDT, ADP, and DJR wrote the manuscript

## 7 Disclaimer

The authors declare no competing interests

Certain commercial equipment, instruments, or materials are identified to adequately specify experimental procedures. Such identification neither implies recommendation nor endorsement by the National Institute of Standards and Technology nor that the equipment, instruments, or materials identified are necessarily the best for the purpose.

