## Supplemental materials for "Interpretable modeling of genotype-phenotype landscapes with state-of-the-art predictive power"

### S1 LANTERN scales to any dataset size

As an additional validation of LANTERN's ability to provide useful insight from GPL data, we assessed its performance on five datasets with 5—7 total mutations (Sailer and Harms 2017). Each dataset was fit with our described approach, using all available measurements. These datasets ostensibly measure the same structure as large-scale GPL datasets, but due to their relative small size they are not amenable to modeling with data-driven methods like deep neural networks. LANTERN, however, is capable of explicitly balancing the amount of data available against the potential complexity of the underlying GPL. LANTERN is therefore capable of learning on these data in exactly the same fashion as higher-throughput measurements (Supplemental Fig S7A-E). In all cases, LANTERN finds a single dimension of relevant variation for these dataset (Supplemental Fig S7F). This single dimension generally captures a saturating non-linear curve as a function of mutational shifts in the latent phenotype space. LANTERN is able to recover interpretable conclusions from these datasets, like the relatively negligible impact of a glycine substitution on transcription factor binding affinity (Fig S7E, Anderson et al. 2015).

### S2 LANTERN generalizes simpler models

LANTERN outperforms simpler GPL models in predictive accuracy (Fig 3). To determine whether there are any similarities between LANTERN and these alternatives, we compared components learned by LANTERN to the simpler approximations. We focused on $\mathbf{z}_1$ because it has the highest variance of any dimension, and therefore contains the most information on the GPL. $\mathbf{z}_1$ correlates strongly both with the average effect of mutations across all backgrounds ($0.58 \leq \rho \leq 0.91$, $p < 10^{-3}$, Fig S12) and the effects learned by a monotonic spline model ($0.660 \leq \rho \leq 0.960$, $p < 10^{-3}$, Fig S12). Additionally, the surface $f(z)$ along $\mathbf{z}_1$ partially resembles the one-dimensional spline model (Supplemental Figs S12, S13). Overall this suggests that the model learned by LANTERN is related to simpler models, but LANTERN extends these approaches to better explain complex landscapes.

### S3 Deriving posterior predictive distrubtion of differential operators

Analytic posterior predictive distributions of differential operators on $f(z)$ were calculated using techniques from Solak et al. 2003. Throughout we describe calculations with respect to the approximate variational posterior $q(f(Z)) = N(m, S)$ at inducing points $Z$. Calculations were simplified by deriving equations for prediction of differential operators for each point individually. We refer to the point where predictions are being made as $z_k$ in the following equations. While derived for predicting an individual point, the same equations are used to predict differential operators at every location of interest on the latent mutational effect space (Fig 6).

The predictive distribution for the gradient $\nabla f(z) = \{\frac{\partial}{\partial z^{(1)}} f(z), \ldots, \frac{\partial}{\partial z^{(L)}} f(z)\}$, where $z^{(i)}$ is the $i$th component of $z$, was computed as

$$q(\nabla f(z_k)) = N(\mu_\nabla, \Sigma_\nabla) \tag{S1}$$

$$= N(K_\nabla K^{-1} m, K_{\nabla\nabla} - K_\nabla K^{-1}(S - K)K^{-1}K_\nabla^T), \tag{S2}$$

where

$$(K_\nabla)_{ij} = Cov\left[\frac{\partial f(z_k)}{\partial z^{(i)}}, f(z_j)\right] \tag{S3}$$

$$= \frac{\partial}{\partial z^{(i)}} Cov\left[f(z_k), f(z_j)\right] \tag{S4}$$

and

$$(K_{\nabla\nabla})_{ij} = Cov\left[\frac{\partial f(z_k)}{\partial z^{(i)}}, \frac{\partial f(z_k)}{\partial z^{(j)}}\right] \tag{S5}$$

$$= \frac{\partial^2}{\partial z^{(i)} \partial z^{(j)}} Cov\left[f(z_k), f(z_k)\right]. \tag{S6}$$

Similarly for the Laplacian $\Delta f(z) = \sum_{i=1}^{L} \frac{\partial^2}{\partial z^{(i)2}} f(z)$, the joint predictive distribution of the individual partial second derivatives were calculated as

$$q\Big(\{\frac{\partial^2}{\partial z^{(i)2}}f(z)|i=1,2,\ldots,L\}\Big) = N(K_\Delta K^{-1}m, K_{\Delta\Delta} - K_\Delta K^{-1}(S-K)K^{-1}K_\Delta^T), \tag{S7}$$

where

$$(K_\Delta)_{ij} = Cov\Bigg[\frac{\partial^2 f(z_k)}{\partial z^{(i)^2}}, f(z_j)\Bigg] \tag{S8}$$

$$= \frac{\partial^2}{\partial z^{(i)^2}} Cov\Bigg[f(z_k), f(z_j)\Bigg], \tag{S9}$$

and

$$(K_{\Delta\Delta})_{ij} = Cov\Bigg[\frac{\partial^2 f(z_k)}{\partial z^{(i)^2}}, \frac{\partial f(z_k)}{\partial z^{(j)}}\Bigg] \tag{S10}$$

$$= \frac{\partial^2}{\partial z^{(i)^2}} Cov\Bigg[f(z_k), f(z_k)\Bigg]. \tag{S11}$$

The predictive distribution of $\Delta f(z) = \mathbf{1}_L^T\{\frac{\partial^2}{\partial z^{(i)2}}f(z)|i=1,2,\ldots,L\}$, where $\mathbf{1}_L$ is an $L \times 1$ vector of ones and induces the summation over the second partial derivatives needed for the Laplacian, is recovered as

$$q(\Delta f(z)) = N(\mu_\Delta, \sigma_\Delta^2) \tag{S12}$$

$$= N(\mathbf{1}_L^T K_\Delta K^{-1}m, \mathbf{1}_L^T(K_{\Delta\Delta} - K_\Delta K^{-1}(S-K)K^{-1}K_\Delta^T)\mathbf{1}_L). \tag{S13}$$

The resulting $q(\nabla f(z))$ and $q(\Delta f(z))$ were then used to quantify robustness and additivity.

| region | residues |
|---|---|
| Dimer interface | 77-100, 221-226, 250-260, 279-285 |
| DNA-binding domain | 6-58 |
| Ligand-associated | 68-70, 73-76, 79, 125-127, 148-150, 160-161, 188, 191, 193-194, 197, 220, 245-249, 273-274, 276, 291, 293, 296, |

**Table S1:** LacI protein structural domains

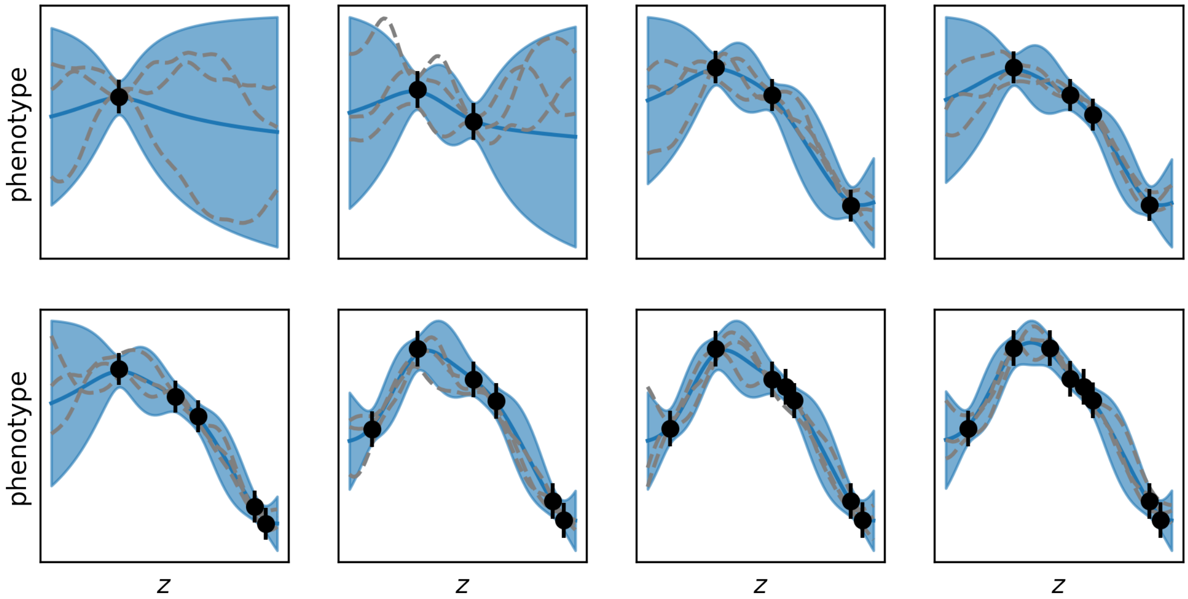


**Figure S1: Sequential inference with Gaussian processes.** Each panel shows the Gaussian process (GP) posterior as noisy, sequential observations are made of the underlying surface $f(\mathbf{z})$: $y(\boldsymbol{z}) = f(\boldsymbol{z}) + \epsilon$ with $\epsilon \sim N(0, \sigma_y^2)$. The posterior of $f(\mathbf{z})$ is shown with the posterior mean (blue solid line) and 95% credible interval (blue shaded region). Individual function draws from the posterior are shown as dotted lines. The available measurements at each step are shown as scatter points with uncertainty as error bars ($2\sigma_y$). At each iteration, the GP posterior balances the experimental evidence against the plausible functions that could have generated the data. As more observations become available, the posterior concentrates around an increasingly certain distribution of these functions. In this way, GPs learn a distribution over functions that most likely generated the data, automatically from the data itself.

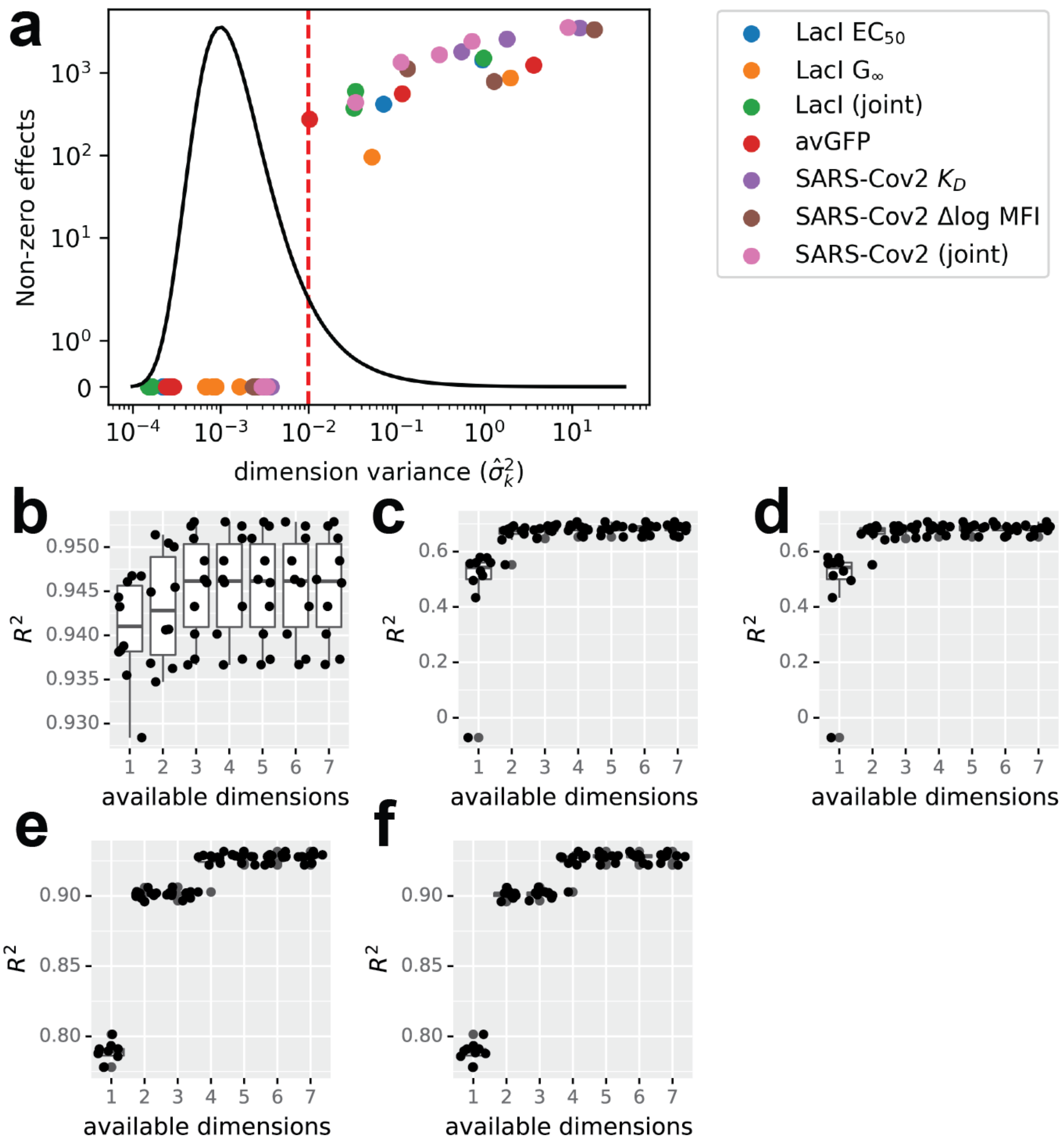


**Figure S2: Determining the dimensionality learned by LANTERN**. (a) The number of statistically significant non-zero effects (at the 95% level) for individual dimensions as a function of learned posterior mean. There is a clear threshold near $\sigma_k^2 = 10^{-2}$. (b-f) Ten-fold cross validation as a function of latent dimensions. Cross-validation predictive accuracy ($R^2$) for LANTERN models with increasing numbers of latent dimensions for datasets: avGFP (a); LacI $EC_{50}$ (b) and $G_\infty$ (c); and SARS-Cov2 binding (d) and expression (e)

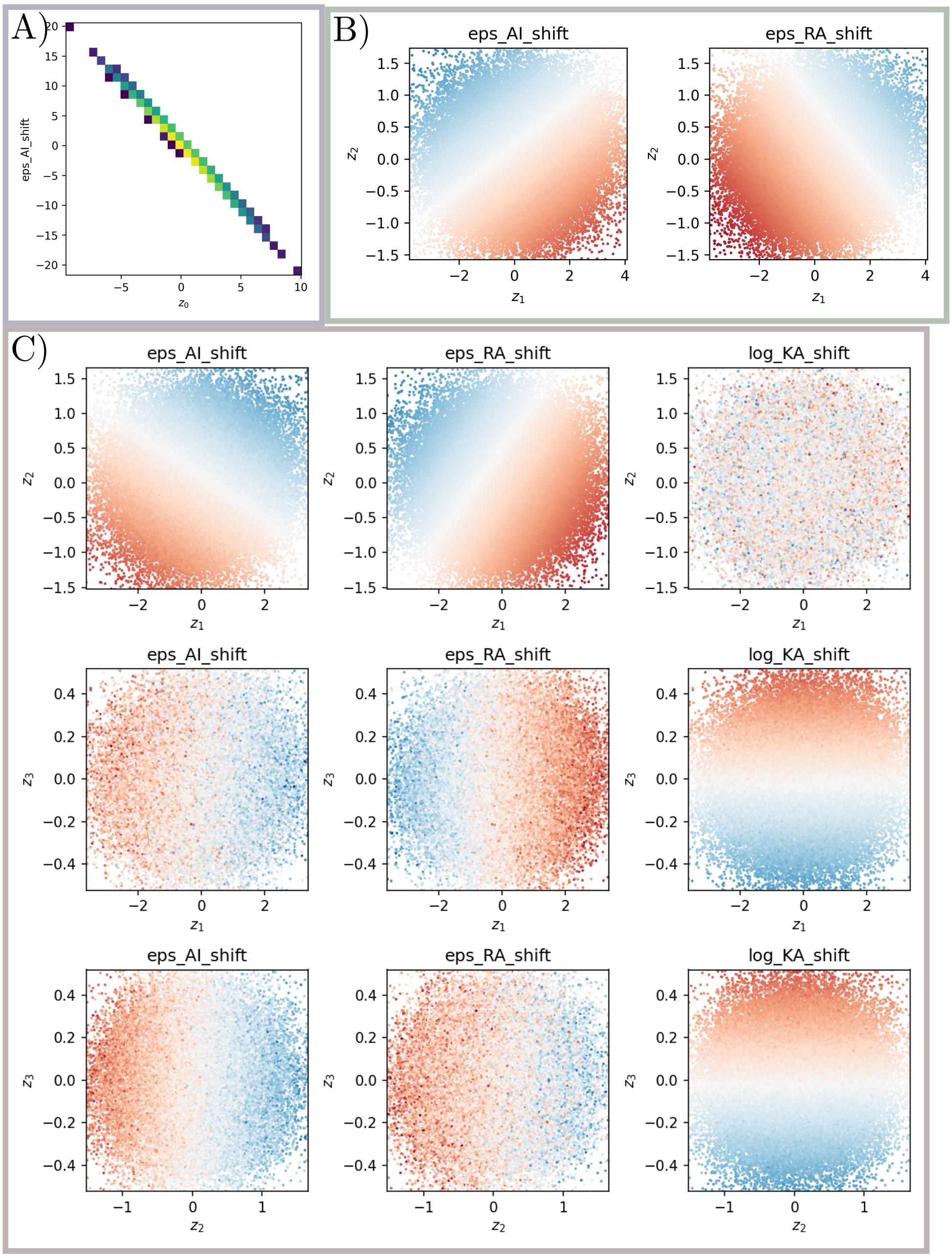


**Figure S3: Correlation between latent dimensions of LANTERN and true biophysical parameters.** (a) Correlation between $\mathbf{z}_1$ and $\Delta\epsilon_{\mathrm{AI}}$in the one-dimensional dataset. (b) Distribution of $\Delta\epsilon_{\mathrm{AI}}$ and $\Delta\epsilon_{\mathrm{RA}}$ along $\mathbf{z}_1$ and $z_2$ in the two-dimensional dataset. Scatter points mark the variant position in $z - space$ and colors are the biophysical parameter values. (c) Distribution of biophysical parameters across different dimensions in the three dimensional dataset. Columns correspond to $\Delta\epsilon_{\mathrm{AI}}$, $\Delta\epsilon_{\mathrm{RA}}$, and $\Delta\log\mathrm{KA}$ respectively and rows show different pairs of $z$ dimensions (1, 2, and 3). Scatter points are variant positions in $z - space$ while colors correspond to the biophysical parameter value. In two and three dimesnions, LANTERN appears to learn rotations of the underlying biophysical parameters $\epsilon_{AI}$ and $\epsilon_{RA}$ rather than partitioning them into independent dimensions. The third dimension, however, was reserved for $\log K_A$, possibly indicating that specifically $\epsilon_{AI}$ and $\epsilon_{RA}$ are indistinguishable from the rotation identified by LANTERN.

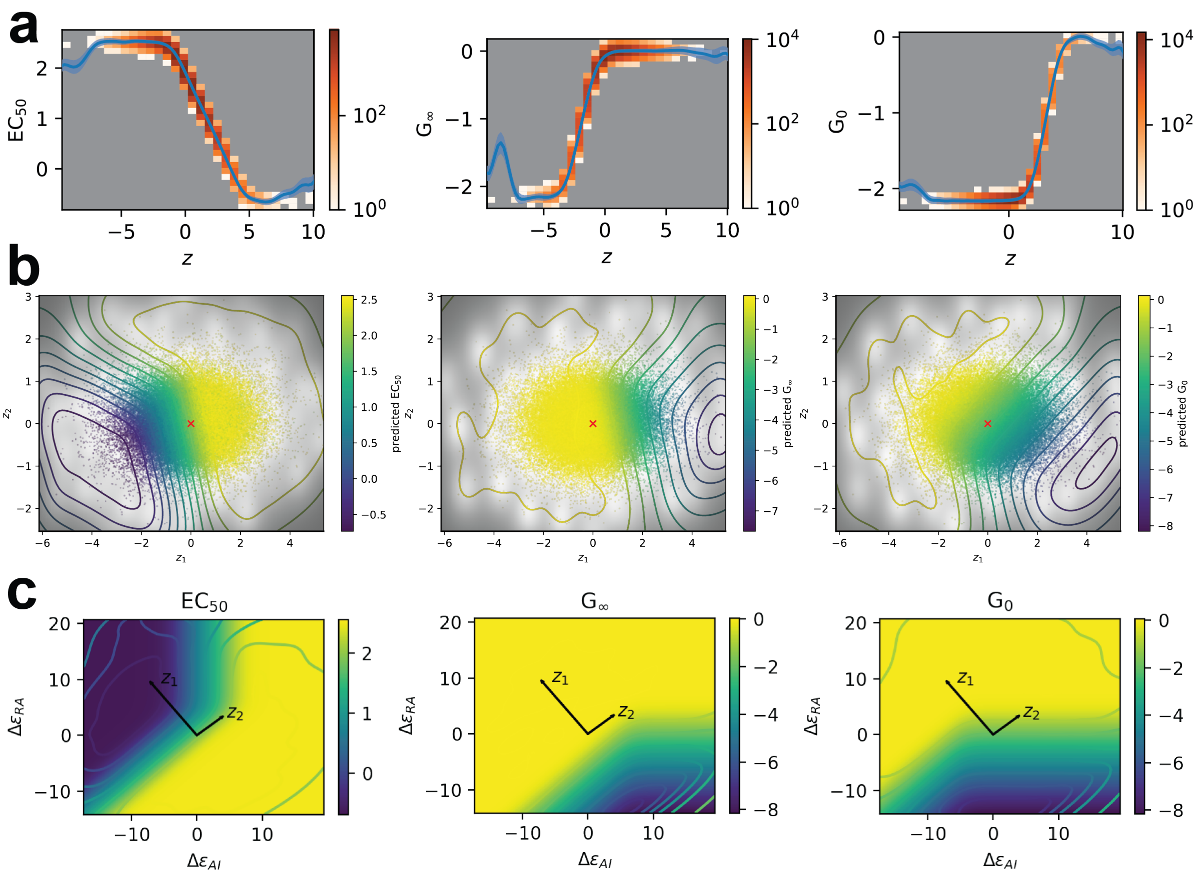


**Figure S4: Recovering biophysical landscapes and dimensions.** (a) LANTERN model for the one-dimensional dataset. Blue line is the posterior mean of $f(z)$ and the shaded region is the 95% credible region. Histograms mark the distribution of observations. (b) Learned two-dimensional surface $f(z)$ for the two dimensional simulation. Contours show the posterior mean of $f(z)$, shading shows the relative variance of $f(z)$ and scatter points are observations positioned by their latent $z$ value and colored by their observed parameter value. (c) Rotation of the learned surface $f(z)$ to the true biophysical surface. As a function of biophysical parameters $\Delta\epsilon_{AI}$ and $\Delta\epsilon_{RA}$, the image shows the true biophysical landscape and the contours show the posterior mean of $f(z)$ from LANTERN. The black vectors mark the rotation of the $\mathbf{z}_1$ and $z_2$ dimensions to the native biophysical space.

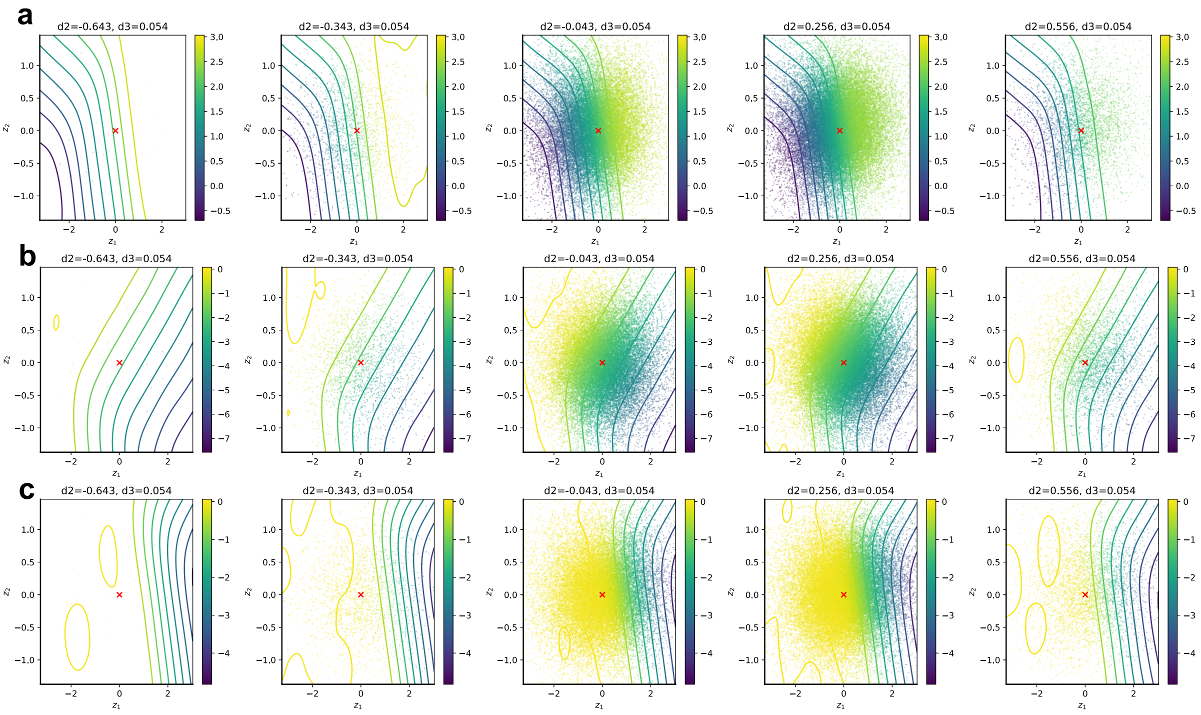


**Figure S5: Scan of $z_3$ of the simulated allostery data**. Scans of the $f(z)$ surface along the third dimension learned by LANTERN for the three dimensional biophysical simulation. Each plot shows the $f(z)$ surface as a function of $\mathbf{z}_1$ and $z_2$ with posterior mean as contours and latent variant positions as scatter points colored by their observed value. Columns correspond to different values of $z_3$ and rows are the surfaces of $EC_{50}$ (a), $G_\infty$ (b) and $G_0$ (c) respectively.

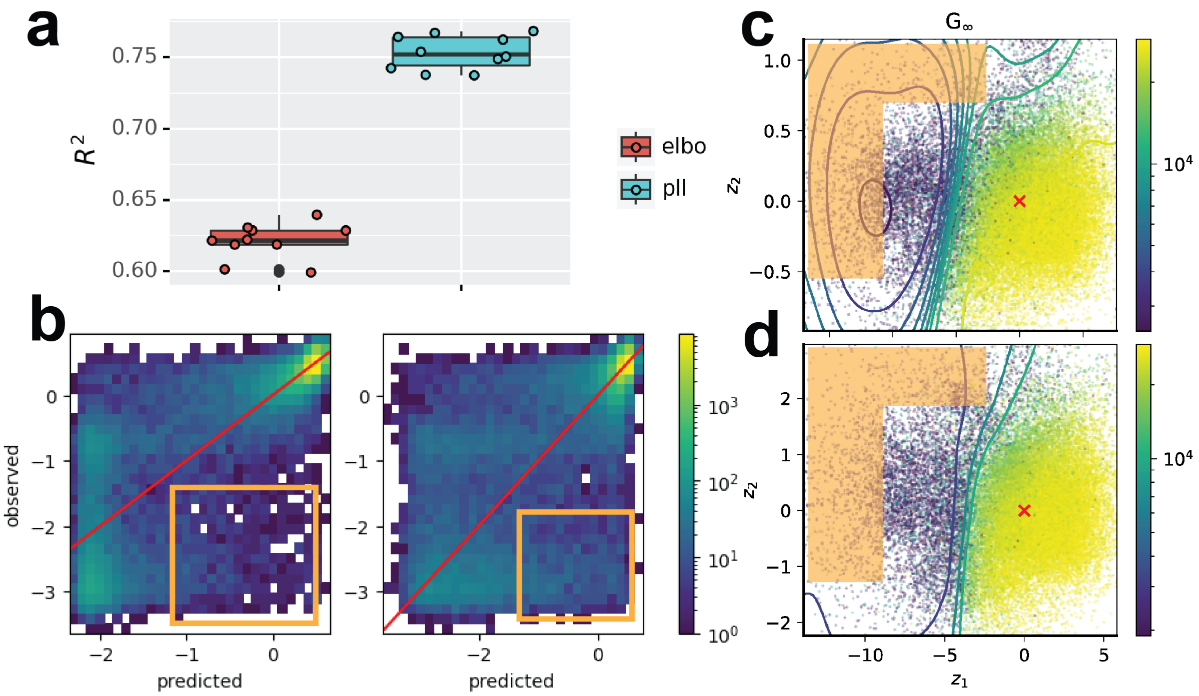


**Figure S6: Improved $G_\infty$ prediction with the predictive log-likelihood objective.** We changed the objective function for the surface $f(\mathbf{z})$ from the usual evidence lower bound (ELBO, Eq 12) to an alternative objective that performs better with large measurement uncertainties called the predictive log-likelihood (PLL) (Jankowiak et al. 2019). (a) When LANTERN is trained with the PLL objective, the cross-validated $R^2$ predictive accuracy for $G_\infty$ increases dramatically. (b) Inspection of model predictions shows that a substantial proportion of the increase in $R^2$ derives from changes in how well LANTERN accurately predicts values of $G_\infty$ that are less than the wild-type when trained with the PLL objective (orange boxes): low predicted values of $G_\infty$ are more likely to be low in reality. This is further demonstrated when reviewing the surfaces learned under an ELBO loss (c) and a PLL loss (d). In the case of the PLL loss, more distant regions of the latent mutational effect space (orange boxes) are more accurately predicted to have $G_\infty$ values different from wild-type. The ELBO trained model (c) displays the more typical Gaussian process behavior of reverting to predicting the mean value in regions with less observations.

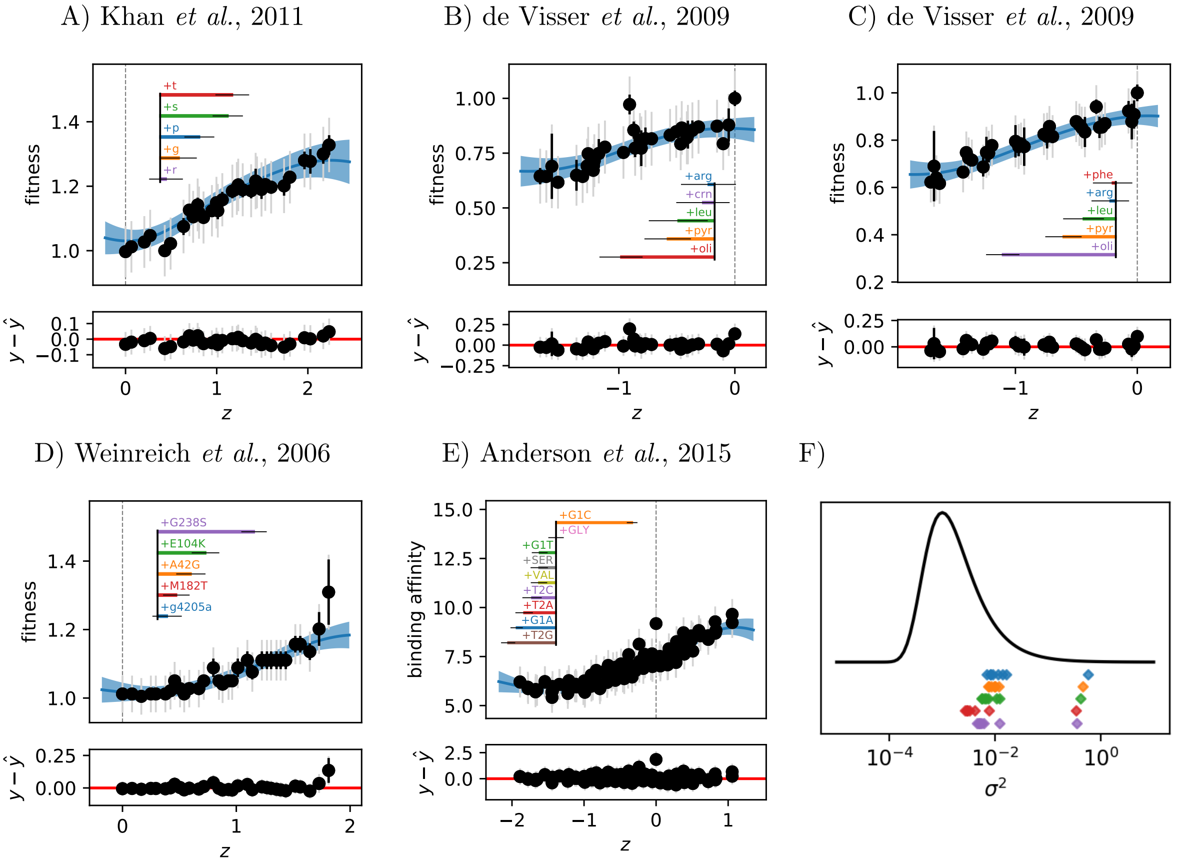


**Figure S7: Interpretable modeling on small scale landscapes.** (A-E) Results of inference on small scale genotype-phenotype landscape datasets. Posterior distribution of $f(z)$ is shown in blue, solid line indicating posterior mean and shaded region 95% credible region. Points show measured values, with the $z$ position set to the posterior mean, black error bars represent reported uncertainty, and grey bars represent learned uncertainty (reported and model uncertainty combined). Posterior mean of mutation effects are shown as colored bars and error bars show 95% intervals. Horizontal line marks the origin. Residuals as a function of $z$ are shown below each model. (F) Posterior variance means for each dataset. Only one dimension has relevant variance in each dataset.

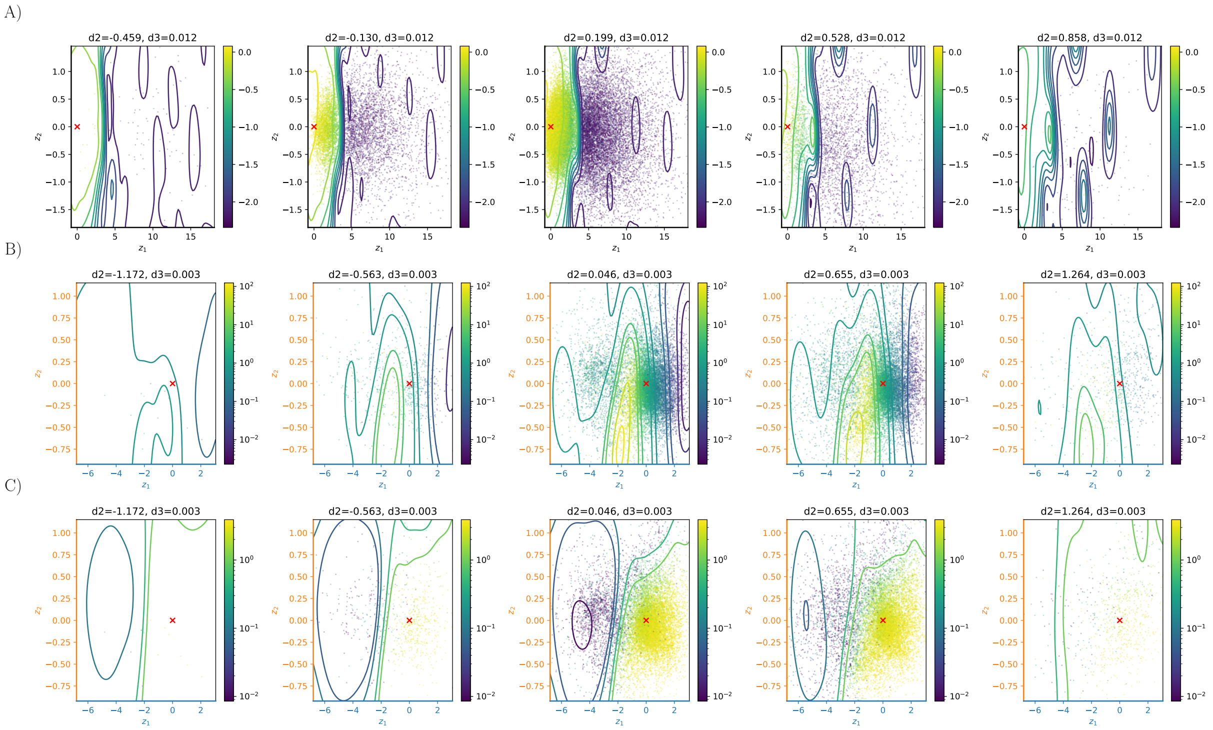


**Figure S8: Higher dimensional scans of avGFP and LacI surfaces.** (A) Scan of $z_3$ dimension of avGFP. (B) Scan of $z_3$ dimension of LacI $EC_{50}$. (C) Scan of $z_3$ dimension of LacI $G_\infty$ Each scan shows the the predicted two dimensional surface $f(z)$ along $\mathbf{z}_1$ and $z_2$, with different values of $z_3$ aranged along each column. Contours show the posterior mean of $f(z)$, scatter points are variant positions in $z$-space colored by their observed value.

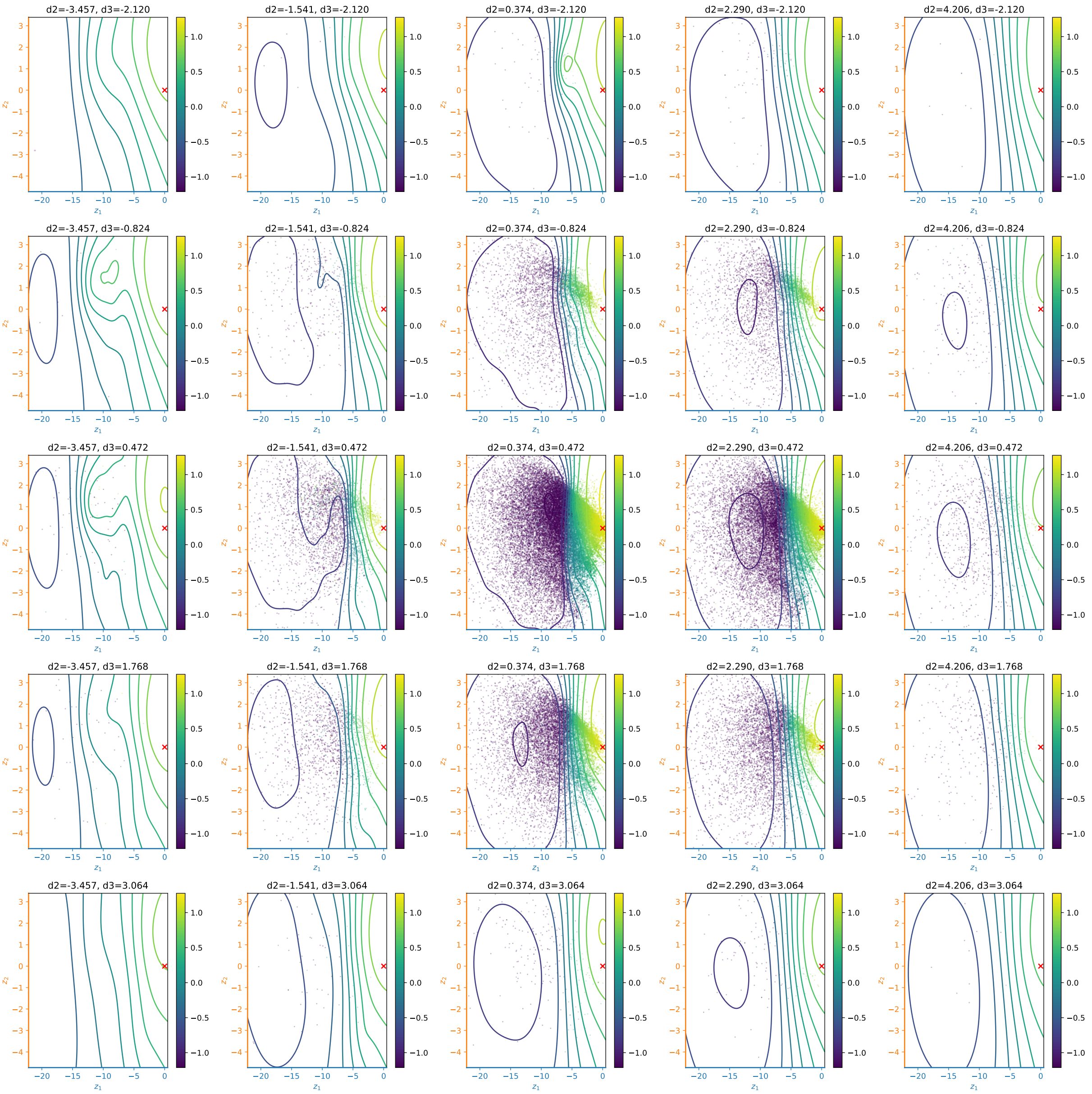


**Figure S9: Scan of higher dimensions of SARS-Cov2 binding surface**. $z_3$ and $z_4$ scan of SARS-Cov2 binding surface. The scan shows the the predicted two dimensional surface $f(z)$ along $\mathbf{z}_1$ and $z_2$, with different values of $z_3$ and $z_4$ arranged along each column and row, respectively. Contours show the posterior mean of $f(z)$, scatter points are variant positions in $z$-space colored by their observed value.

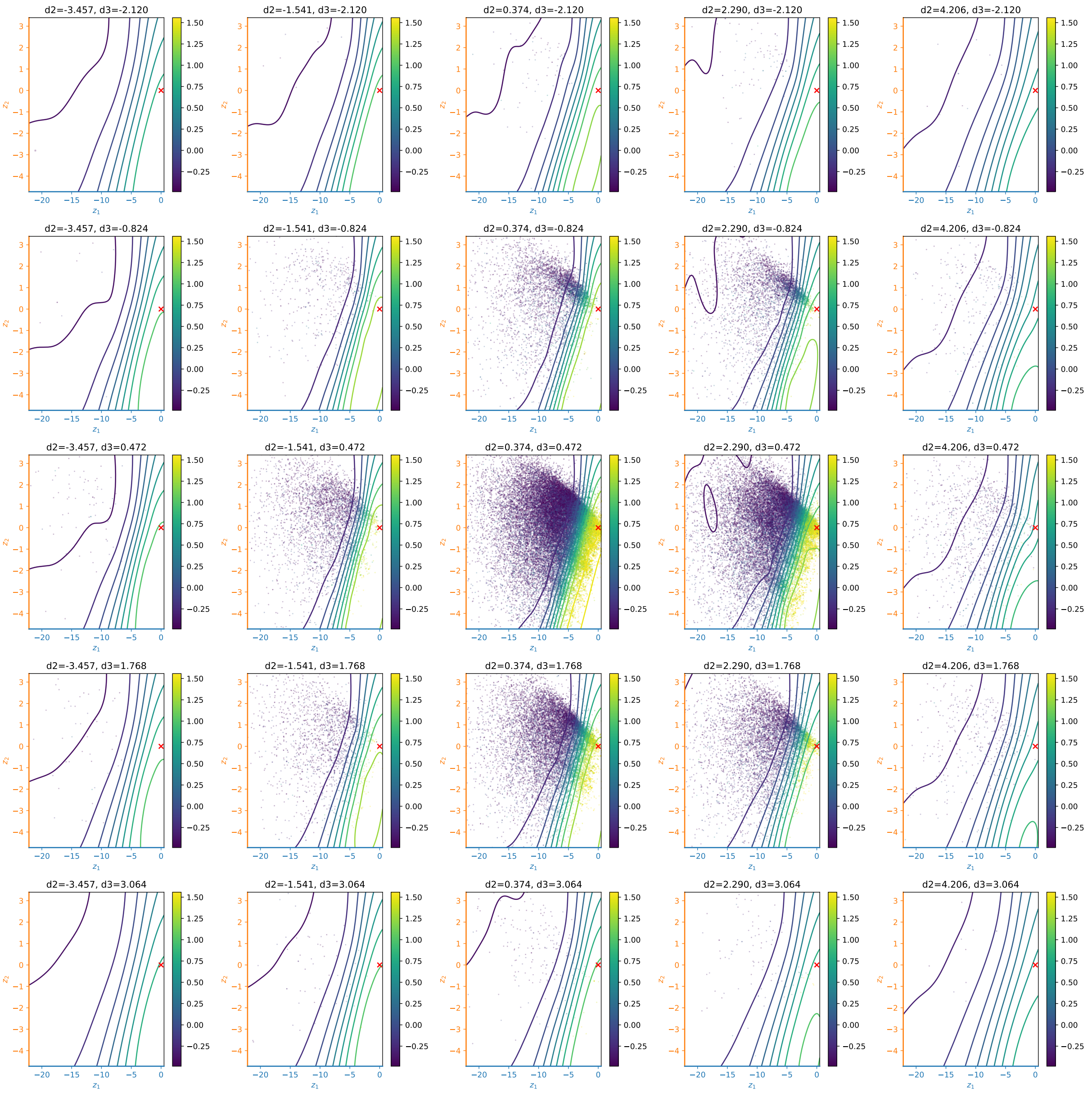


**Figure S10: Scan of higher dimensions of SARS-Cov2 expression surface**. $z_3$ and $z_4$ scan of SARS-Cov2 expression surface. The scan shows the the predicted two dimensional surface $f(z)$ along $\mathbf{z}_1$ and $z_2$, with different values of $z_3$ and $z_4$ arranged along each column and row, respectively. Contours show the posterior mean of $f(z)$, scatter points are variant positions in $z$-space colored by their observed value.

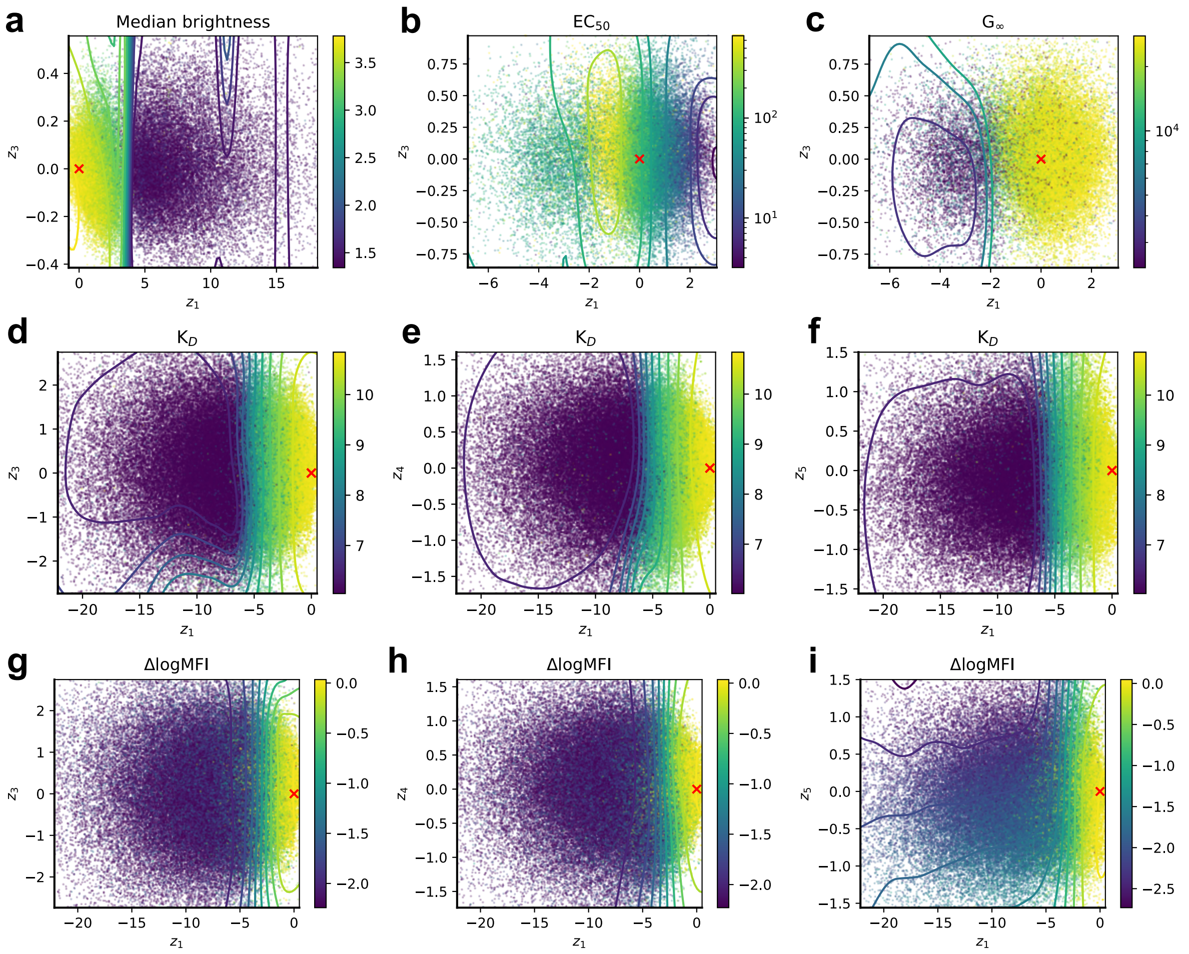


**Figure S11: Surfaces along $\mathbf{z}_1$ versus higher dimensions**. $f(z)$ two-dimensional surfaces along $\mathbf{z}_1$ and higher dimensions $z_3$ — $z_5$. Surfaces show the posterior mean of $f(z)$ as contours with variant distributions in the latent space as scatter points colored by their observed value. Surfaces correspond to avGFP brightness (a), LacI $EC_{50}$ (b) and $G_\infty$ (c), and SARS-Cov2 binding (d—f) and expression (g–i).

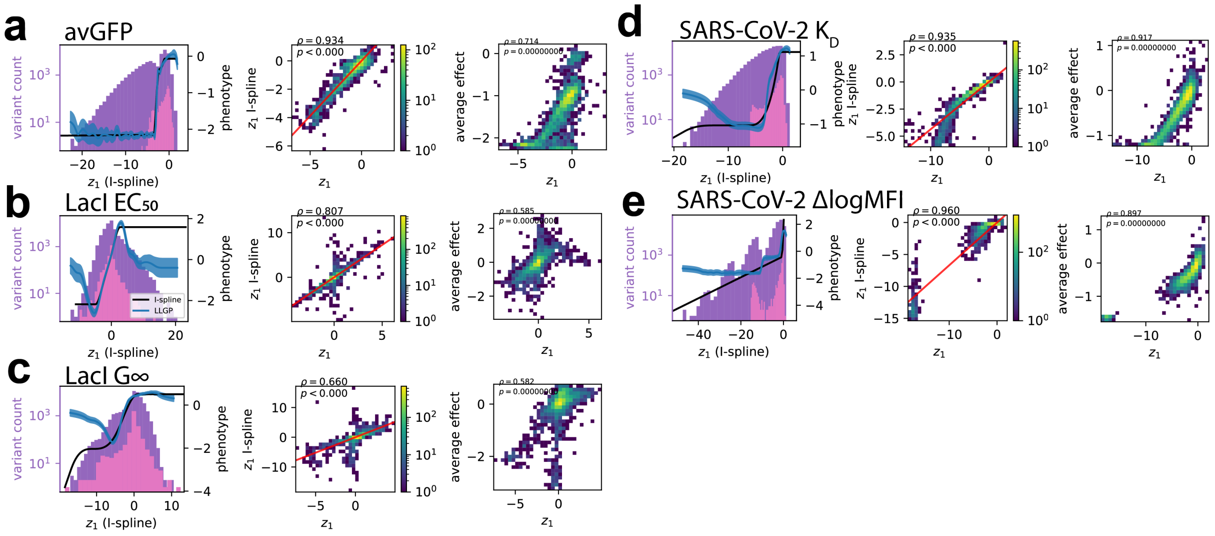


**Figure S12: Comparison of $\mathbf{z}_1$ to alternative one-dimensional models**. We compare the learned $\mathbf{z}_1$ dimension from LANTERN to the single dimension of an I-spline global epistasis model (left, center) and the average effects of each mutation (right) across the large-scale datasets: avGFP brightness (a), LacI $EC_{50}$ (b), LacI $G_\infty$ (c), SARS-Cov2 binding (d), and SARS-Cov2 expression (e). (left) For each phenotype, the I-spline prediction along the latent dimension is shown in black, with the LANTERN posterior of $f(z)$ along $\mathbf{z}_1$ shown in blue (posterior mean as solid line and shaded region for 95% credible region). The distribution of mutational effects from LANTERN is shown in pink, and the overall distribution of variants along $\mathbf{z}_1$ shown in purple. The correlation between $\mathbf{z}_1$ effects and the I-spline effects is shown on the right. (center) Distribution of mutational effects along $\mathbf{z}_1$ compared to the mutational effects estimated in the one-dimensional I-spline model. The correlation and associated p-value are shown in the top-left corner. (right) Correlation between mutational effects along $\mathbf{z}_1$ and the average effect of a mutation across all backgrounds. Correlation and associated p-value are shown in the top-left corner.

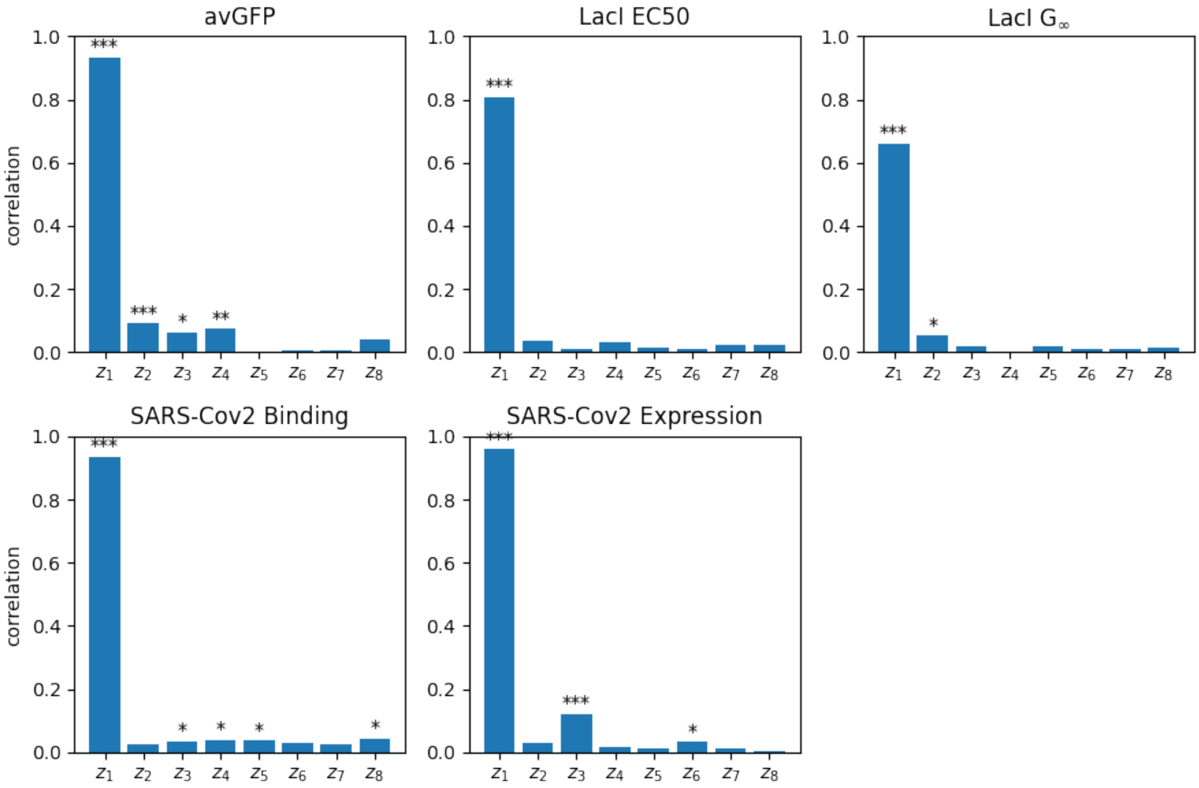


**Figure S13: First dimension correlation with I-spline model**. Correlation between mean latent effect across different dimensions identified by LANTERN and the single latent dimension of the I-spline model from (Jakub Otwinowski et al. 2018). Signficant correlations are marked with p-values less than 0.05 (*), 0.005 (**) and 0.0005 (***).

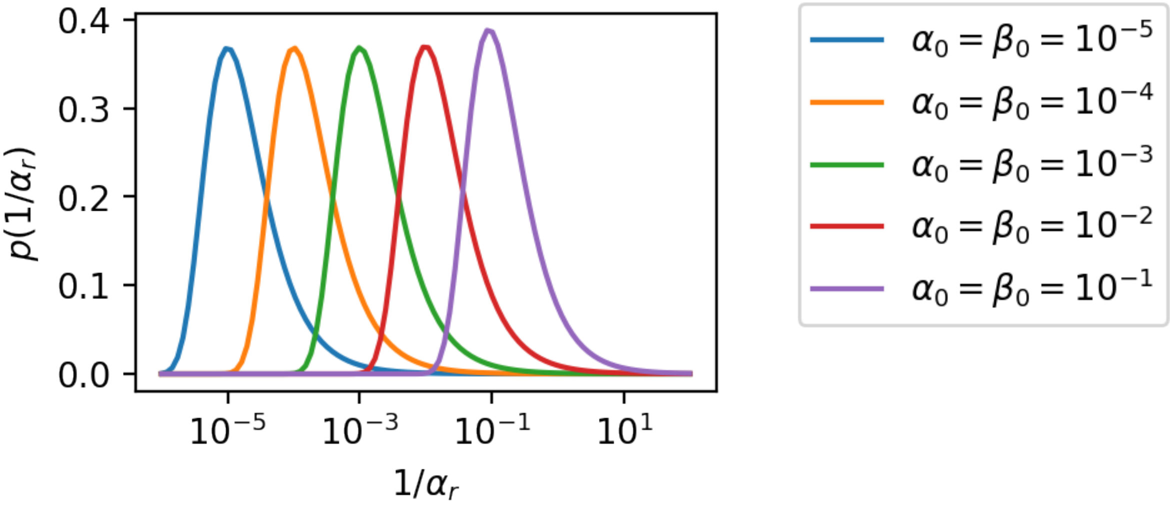


**Figure S14: Effect of changing hyperpriors $\alpha_0$ and $\beta_0$.** For different values of $\alpha_0$ and $\beta_0$, the resulting prior on the variance $(1/\alpha_r)$ for each latent dimension $r$ in the model is shown. See Equation 5.

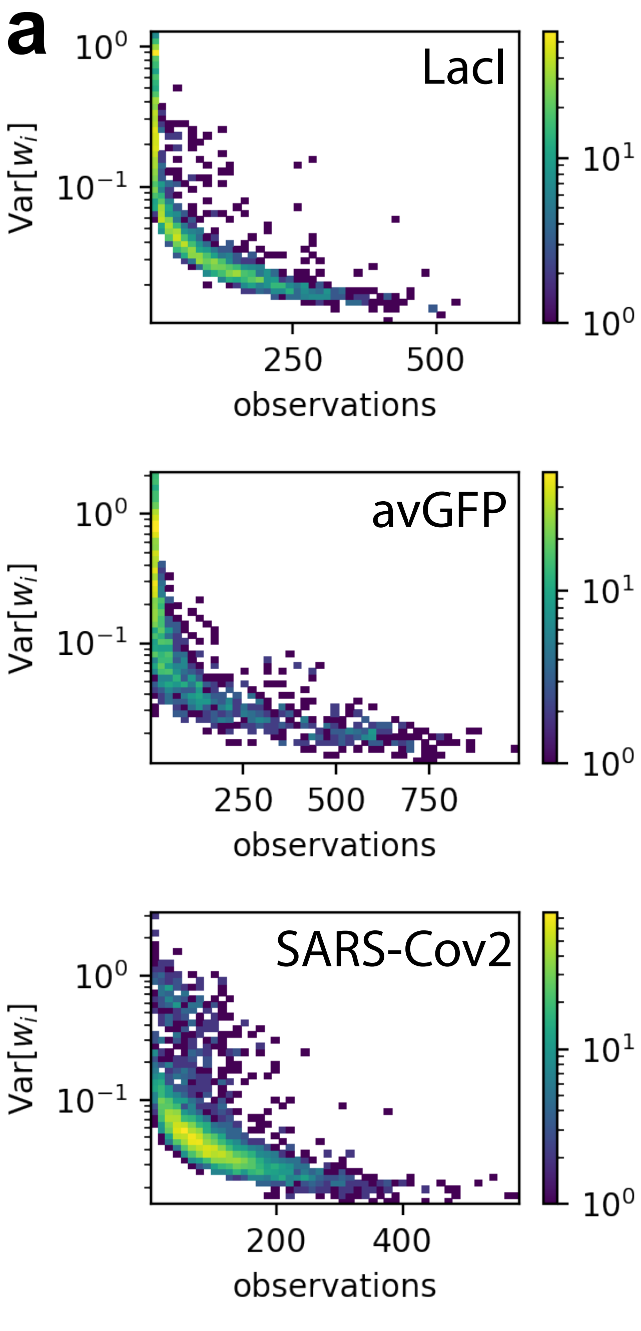


**Figure S15: Uncertainty of LANTERN components**. (a) Posterior variance of mutational effects as a function of unique variants containing the corresponding mutation.

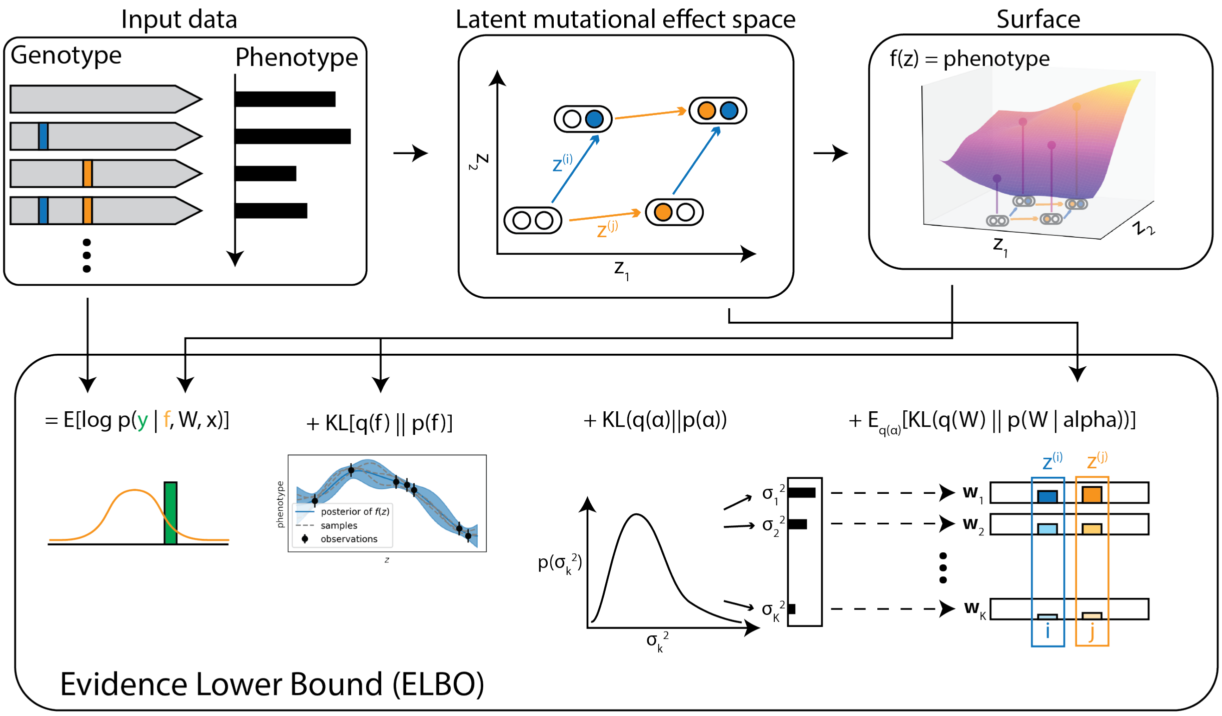


**Figure S16: Calculating the evidence lower bound (ELBO) for LANTERN.** The ELBO is computed as a combination of the expected log probability of observations and the KL-divergence of the approximated posterior for each model component from its prior.

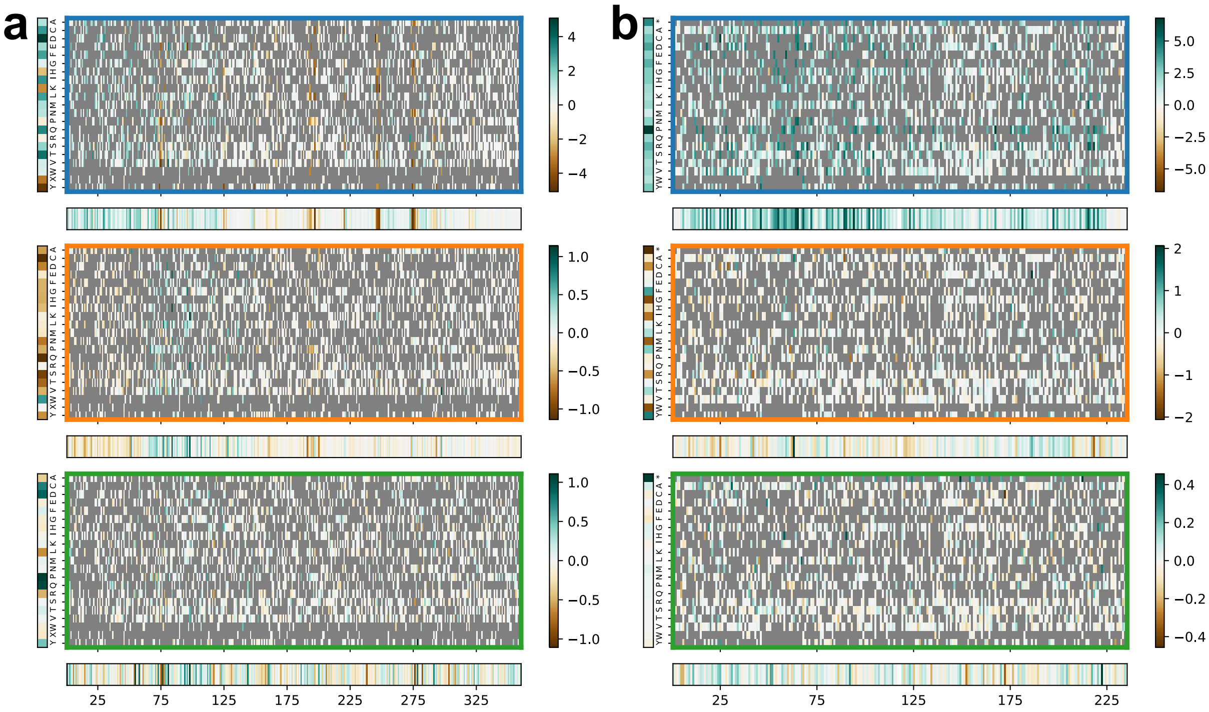


**Figure S17: Mutation effect scores of LacI and GFP**. Posterior mean effects for each mutation across $\mathbf{z}_1$, $z_2$, and $z_3$ are shown for the LacI (a) and avGFP models (b). Unobserved mutations are marked in grey. Average amino-acid effects and average position effects are shown to the left and below each heatmap, respectively.

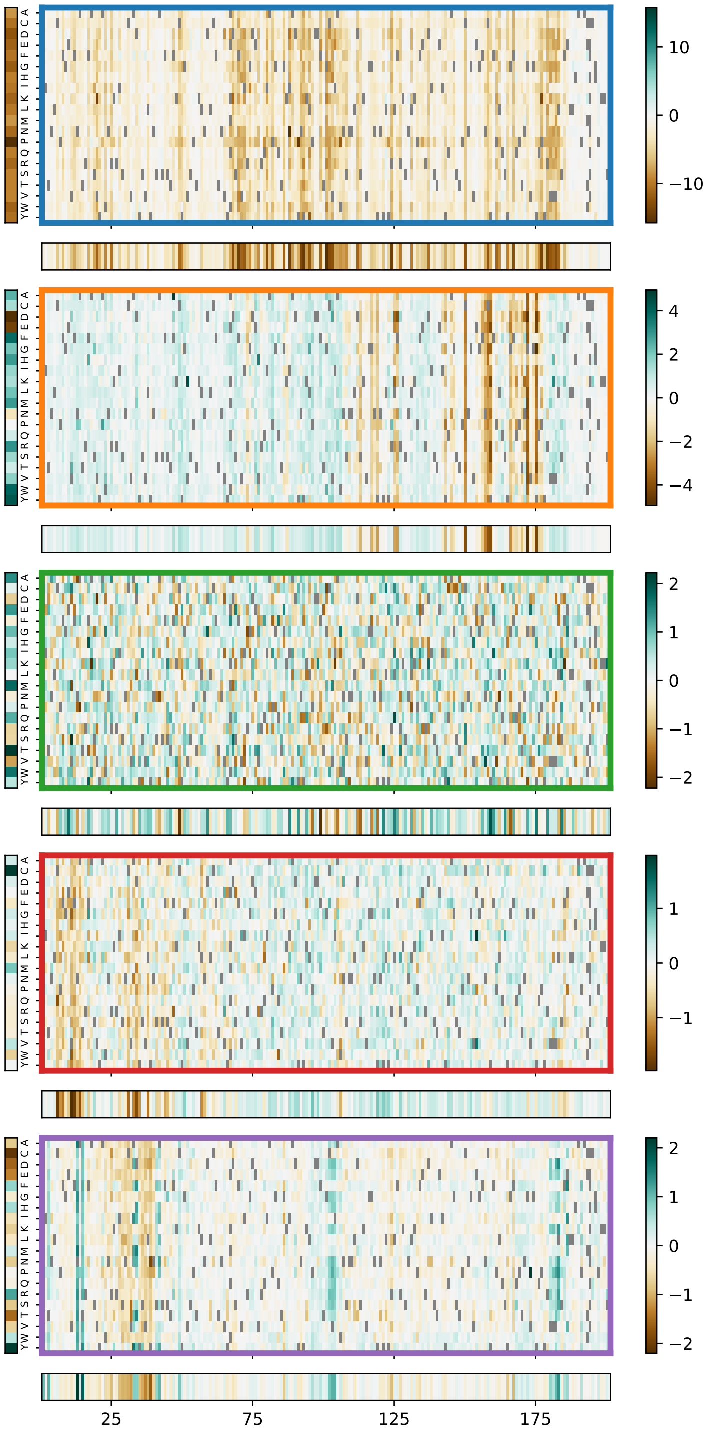


**Figure S18: Mutation effect scores of SARS-Cov2**. Posterior mean effects for each mutation across the five dimensions of the SARS-Cov2 model. Unobserved mutations are marked in grey. Average amino-acid effects and average position effects are shown to the left and below each heatmap, respectively.

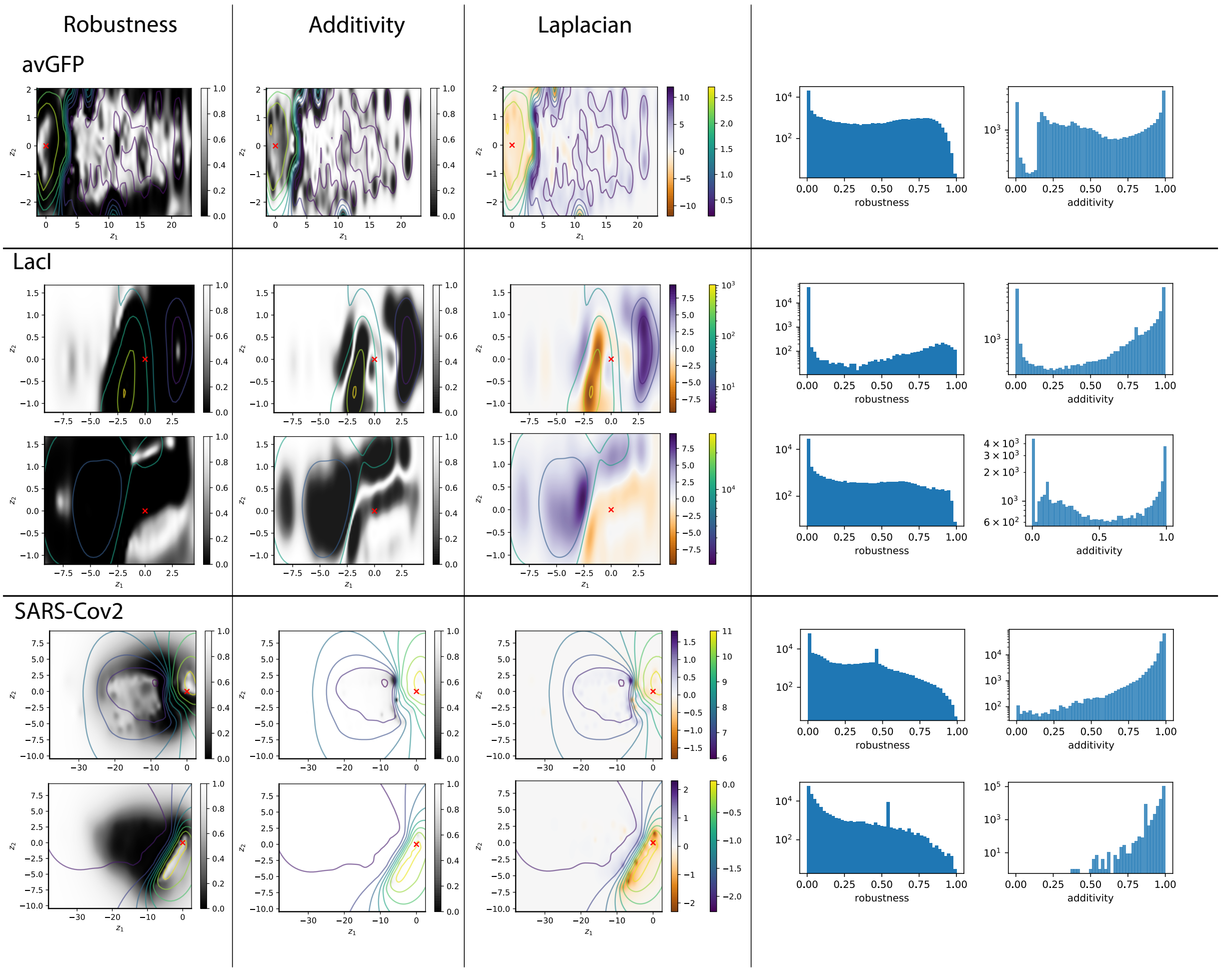


**Figure S19: Differential operators learned by LANTERN**. Robustness, additivity, and mean Laplacian surfaces are shown for each of avGFP, LacI and SARS-cov2 along $\mathbf{z}_1$ and $z_2$ for each dataset. For robustness and additivity, the metric value is shown as a black-to-white heatmap while the posterior mean of $f(z)$ is shown as contours. For the Laplacian, the mean Laplacian value is shown as a heatmap while the contour marks the posterior mean of $f(z)$. Distribution of robustness and additivity values for observed variants is shown as histograms for each dataset.
